# Capturing the onset of PRC2-mediated repressive domain formation

**DOI:** 10.1101/272989

**Authors:** Ozgur Oksuz, Varun Narendra, Chul-Hwan Lee, Nicolas Descostes, Gary LeRoy, Ramya Raviram, Lili Blumenberg, Kelly Karch, Pedro R. Rocha, Benjamin A. Garcia, Jane A. Skok, Danny Reinberg

## Abstract

Polycomb repressive complex 2 (PRC2) maintains gene silencing by catalyzing methylation of histone H3 at lysine 27 (H3K27me2/3) within chromatin. By designing a system whereby PRC2-mediated repressive domains were collapsed and then reconstructed in an inducible fashion *in vivo*, a two-step mechanism of H3K27me2/3 domain formation became evident. First, PRC2 is stably recruited by the actions of JARID2 and MTF2 to a limited number of spatially interacting “nucleation sites”, creating H3K27me3-forming polycomb foci within the nucleus. Second, PRC2 is allosterically activated via its binding to H3K27me3 and rapidly spreads H3K27me2/3 both in cis and in far-cis via long-range contacts. As PRC2 proceeds further from the nucleation sites, its stability on chromatin decreases such that domains of H3K27me3 remain proximal, and those of H3K27me2 distal, to the nucleation sites. This study demonstrates the principles of *de novo* establishment of PRC2-mediated repressive domains across the genome.

## Introduction

Cellular diversity during development arises from the formation of stable and heritable repertoires of gene expression profiles, unique to individual lineages. Genetic and biochemical studies have demonstrated that some histone post-translational modifications (hPTMs) are critical determinants to establishing and maintaining cellular identity (Bonasio et al., 2010; Margueron and Reinberg, 2010). Amongst them, di-and tri-methylation of histone H3 at lysine 27 (H3K27me2/3) – modifications catalyzed by Polycomb repressive complex 2 (PRC2) – are associated with a repressive chromatin state (Ferrari et al., 2014; Lee et al., 2015; Margueron and Reinberg, 2011a; Pengelly et al., 2013; Simon and Kingston, 2013). PRC2 can bind to the products of its own catalysis, H3K27me2/3, through an aromatic cage present in its EED subunit (Margueron et al., 2009). In particular, binding to H3K27me3 leads to an allosteric activation of its methyltransferase activity. We proposed a model whereby the initial catalysis of H3K27me3 at specific sequences could serve to stimulate further H3K27me3 catalysis via a “read-write” mechanism to form large repressive domains as found, for example, across the *Hox* clusters (Boyer et al., 2006).

While there is a large body of data focusing on polycomb genomic localization in mammals, the mechanism(s) by which PRC2 is initially recruited to specific genomic loci and how PRC2 sets up repressive domains comprising H3K27me2 or H3K27me3 remain unclear. We and others, discovered that the genomic targeting of PRC2 is facilitated at developmental genes by its interacting partner JARID2 (Landeira et al., 2010; Li et al., 2010; Pasini et al., 2010; Peng et al., 2009; Shen et al., 2009), a protein that can bind directly to nucleosomes (Son et al., 2013) and exhibits a low affinity for GC-rich sequences *in vitro* (Li et al., 2010). In addition, a recent study demonstrated that MTF2, a member of the Polycomb-like (PCL) proteins that forms a PRC2 sub-complex excluding JARID2 (Grijzenhout et al., 2016), is required for efficient recruitment of PRC2 to CpG islands in mESCs (Li et al., 2017). Both of these studies, as well as others (for review see Holoch et al., 2017 (Holoch and Margueron, 2017)), were performed in mESCs under steady-state conditions, which are not conducive to tracking the *de novo* establishment of PRC2-mediated repressive domains, including the dynamics of and determinants to initial PRC2 recruitment and the behavior of PRC2 as it progressively builds-up H3K27me2/3 domains.

Thus, we devised a genetic system in mESCs by which we could inducibly reconstruct these domains from scratch. Employing this system, we defined the exact genomic coordinates of PRC2 nucleation and spreading and identified the key factors for its initial recruitment and stability on chromatin.

## Results

### Disruption of EED-H3K27me3 interaction reduced H3K27me3 levels in mESCs

The crystal structure of EED shows that H3K27me3 is located within an aromatic cage formed by the WD40-repeats of EED and that three amino acids, Phe97, Tyr148 and Tyr365 directly contact the tri-methylated lysine residue (Margueron et al., 2009). To test whether this interaction was necessary for the maintenance of repressive chromatin domains *in vivo*, we used the CRISPR/Cas9 system in mESCs to generate EED cage-mutants having Phe97 or Tyr365 substituted with alanine (F97A or Y365A). As a control, we mutated Tyr358 (Y358A), which does not contact the tri-methylated lysine. A western blot (WB) of whole-cell extract preparations from these cage-mutant lines showed a significant reduction in H3K27me3 levels (Figure 1a), while those of the control Y358A mutant were unaffected. The global levels of another repressive mark, H3K9me3, remained unperturbed, suggesting that the effect was specific to H3K27me3 (Figure 1a). Quantitative mass-spectrometry of acid-extracted histones demonstrated a precipitous drop in the percentage of the H3K27me3 modification, from 20% in wild type (WT) ESCs to approximately 1% in each of the cage-mutant backgrounds, compared to 0% in the EED knockout (KO) control (Figure 1b).

EED cage-mutant cells also showed a substantial decrease in H3K27me2 levels (Figure 1b), and a reciprocal increase in the levels of H3K36me2, a modification associated with active transcription (Figure S1a), and mutually exclusive with H3K27me3 (Schmitges et al., 2011; Yuan et al., 2011; Zheng et al., 2012). No other quantifiable histone modifications were significantly altered (Table S1). Importantly, the cage mutation did not disrupt the integrity of the PRC2 complex, in contrast to the EED KO (Figure S1b and c). Furthermore, similar to polycomb mutants (Margueron and Reinberg, 2011b), cage-mutant mESCs exhibited WT morphology and gene expression profiles (Figure S2a, b), but failed to differentiate (Figure S2c, d, and e). These results suggest that the EED-H3K27me3 interaction is dispensable for maintaining the pluripotency of mESCs, but required for *de novo* H3K27me3 repressive chromatin domains during differentiation.

**Figure 1:**
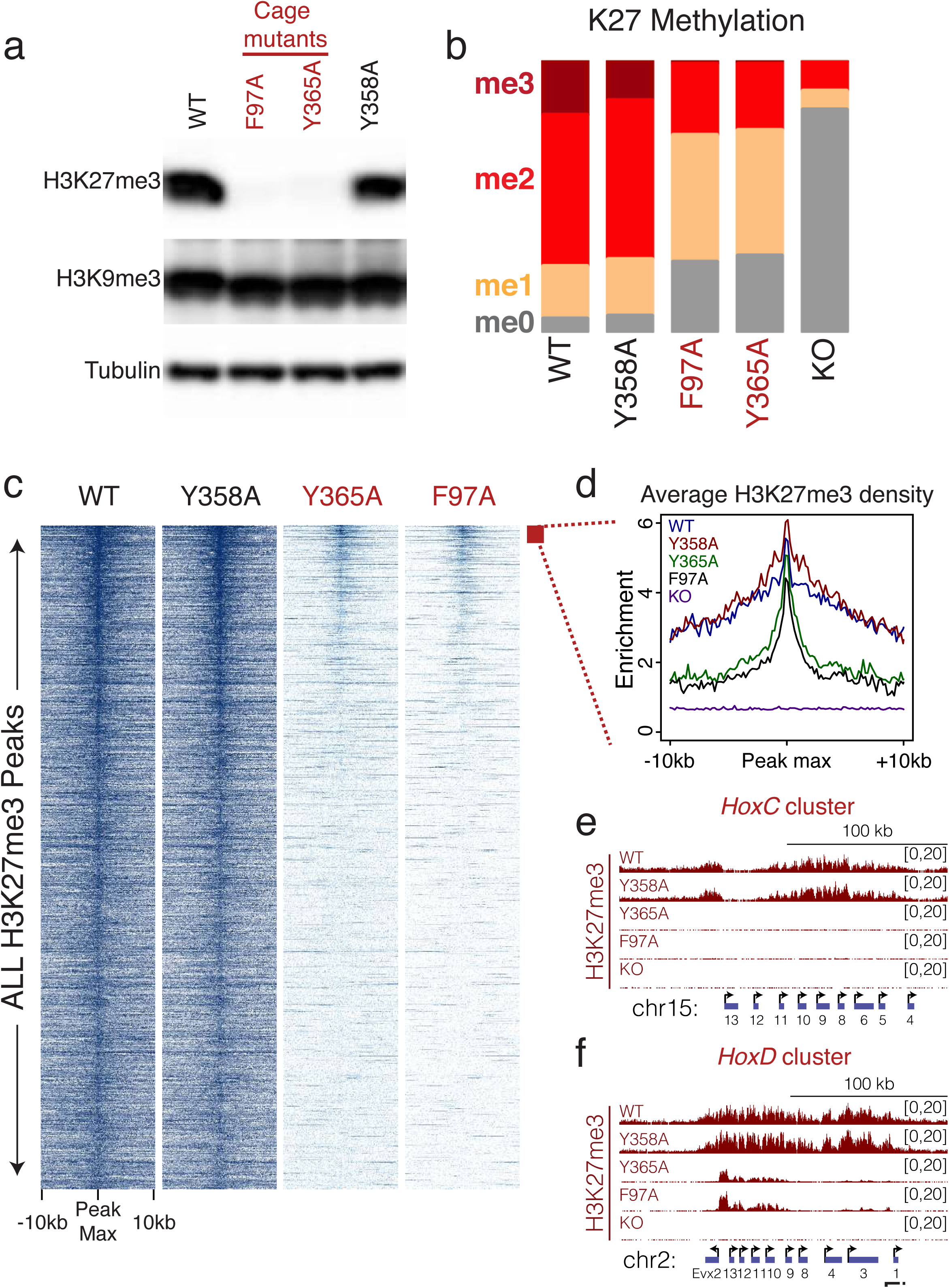
Disruption of the EED-H3K27me3 interaction led to reduced H3K27me3 levels in mESCs. **a,** Western blot using the indicated antibodies on whole cell extracts from E14-TG2a mESCs with the following genotypes of EED; WT: wild type; F97A: cage-mutant; Y365A: cage-mutant; Y358A: control mutant. **b,** Global % of H3K27 methylation levels in mESCs with the indicated genotypes of EED, as identified by quantitative histone mass-spectrometry. **c,** Heatmaps of H3K27me3 ChIP-seq read density within a 20 kb window centered on the maximum value of the peak signal in WT mESCs. H3K27me3 peaks (total=27281 peaks) were sorted by decreasing signal intensity in Y365A cage-mutant mESCs (Scale: 0-1). **d,** Average H3K27me3 density profiles of the top 200 peaks from the indicated genotypes of EED. Of note, EED KO cells have a homozygous frame shift mutation in exon 10, which leads to a premature stop codon and destabilization of the protein. **e, f**, Representative H3K27me3 ChIP-seq tracks at the *HoxC* **(e)** and *HoxD* **(f)** clusters.

Though significantly diminished, H3K27me3 was still detectable in cage-mutant mESCs. To identify the genomic coordinates of the residual ∼1% of H3K27me3, we performed quantitative ChIP-seq (Bonhoure et al., 2014; Egan et al., 2016; Orlando et al., 2014) in WT and cage-mutant cells (Figure 1c). As expected from our global analysis, the cage-mutants showed a significant reduction in H3K27me3 levels across all PRC2 target genes (Figure 1c). For example, H3K27me3 was undetectable across the entire *HoxC* cluster, mimicking the EED KO setting (Figure 1e). Nonetheless, ∼WT levels of H3K27me3 were maintained in cage-mutant mESCs at a limited number of discrete regions within larger polycomb domains, with a progressive decrease distal to these sites (Figure 1d). Of ∼200 such sites, 73% localized to within +/-5 kb of the transcriptional start sites (TSS) of genes. For example, Y365A mESCs displayed WT levels of H3K27me3 at the *Evx2* promoter, yet these levels progressively diminished across the neighboring *HoxD* cluster as a function of the genomic distance from the *Evx2* TSS (Figure 1f). These ChIP-seq results were validated by ChIP-qPCR at a number of target loci (Figure S3). Of note, PRC2 was strictly localized to sites retaining H3K27me3 in the EED cage-mutant mESCs as evidenced by ChIP-seq for EED and SUZ12 (Figure S4e and h (red), respectively). Thus, PRC2 is actively recruited to a discrete number of sites on the genome and its EED-H3K27me3 interaction is required for spreading H3K27me3 in cis along the chromosome.

### Initial deposition and spreading mechanism of H3K27me2/3 domains

To test whether regions of active PRC2 recruitment identified above, might serve as nucleation sites for the initial establishment of polycomb chromatin domains, we devised a system in which new domains could be established “*de novo*” in an inducible fashion *in vivo* and their formation tracked over time. To this end, we deleted the endogenous copy of EED in mESCs possessing a CreERT2 transgene (Figure 2a). These cells did not exhibit H3K27me2/3 deposition as evidenced by WB analysis, and thus provided a demethylated platform upon which to detect *de novo* H3K27me2/me3 deposition (Figure S4a, see 0 hr). We then expressed a FLAG-HA tagged WT or cage-mutant EED protein from its endogenous locus along with a T2A-GFP marker to identify cells in which recombination was successful (Figure 2a, bottom and Figure S4a). These inducible WT or cage-mutant <u>r</u>escue cells were termed “i-WT-r” or “i-MT-r”, respectively.

**Figure 2:**
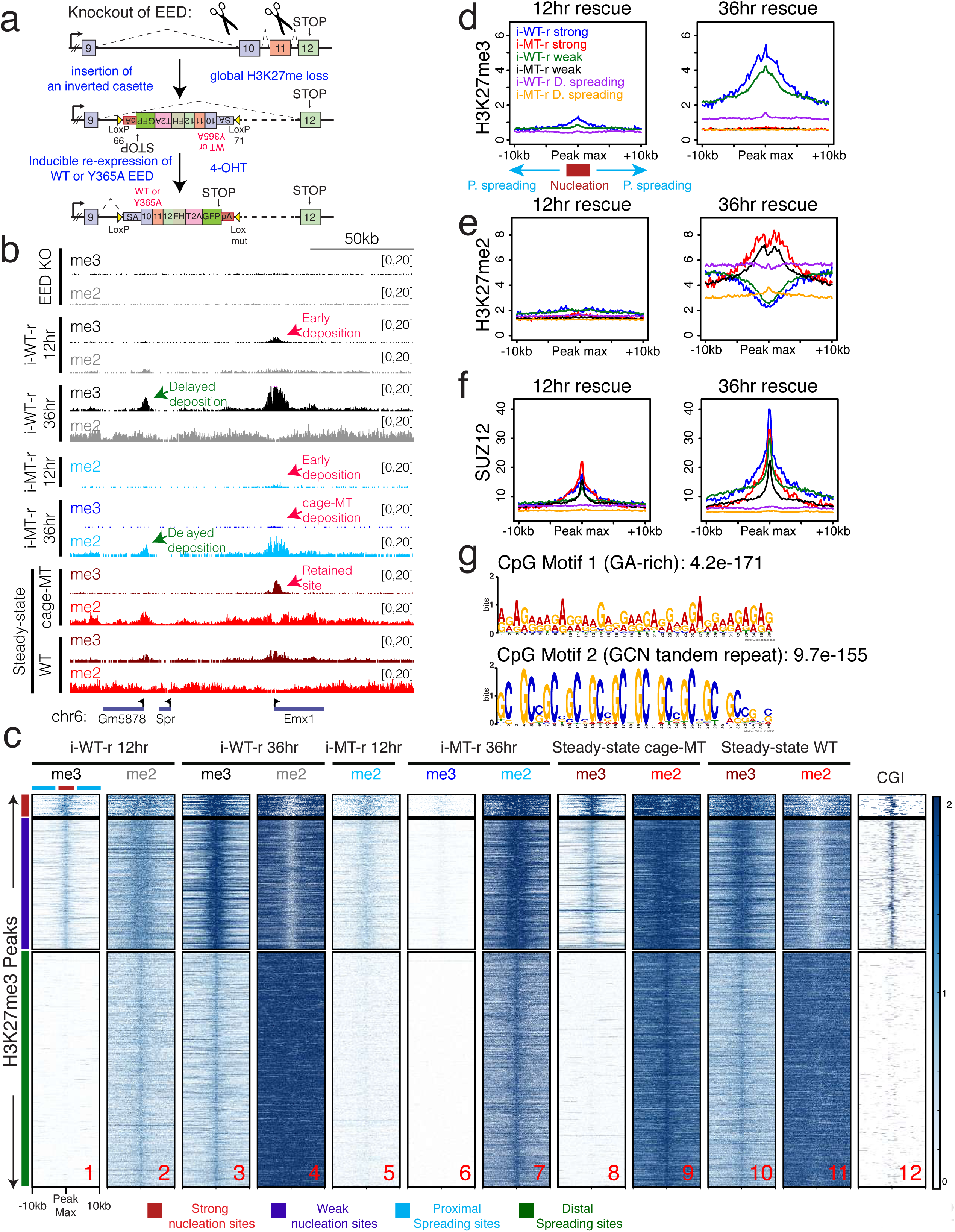
PRC2 activity propagates from initial regions of high CGI density enriched with specific motifs. **a,** Targeting scheme to conditionally rescue EED KO mESCs (C57BL/6) with EED, either WT or cage-mutant (Y365A) (see methods for details). **b,** ChIP-seq tracks for H3K27me3 (me3) and H3K27me2 (me2) near the *Emx1* gene in the EED KO, or in i-WT-r or i-MT-r cells after rescue with EED, either WT or cage-mutant, respectively, for the indicated times of 4-OHT treatment, or in steady-state cells expressing cage-mutant Y365A or WT EED. **c,** Heatmaps of H3K27me3 and H3K27me2 for the rescue experiments performed as in **b**., and CGI density within a 20 kb window centered on the maximum value of peak signal in WT mESCs. H3K27me3 peaks were sorted in descending order by signal intensity at the 12 hr time-point. Strong nucleation sites are marked in red (total=237), weak nucleation sites in blue (total=1389), proximal spreading sites in cyan and distal spreading sites in green (total= 36618, representative of randomly selected 2500 peaks are used for the Heatmaps). (Scales: 0-2 for H3K27me3 and H3K27me2; 0-50 for CGIs). **d, e, f,** Average density profiles of H3K27me3 **(d)** H3K27me2 **(e)** and SUZ12 **(f)** on nucleation sites (strong and weak) and spreading sites (D=distal, P=proximal) derived from the genotypes indicated within a 20 kb window centered on the maximum value of peak signal in WT mESCs. i-WT-r and i-MT-r cells were rescued with EED, either WT or cage-mutant, respectively, for 12 hr (left) or 36 hr (right) of 4-OHT treatment, as indicated. **g,** Motif analysis of strong nucleation site CGIs (∼200 sites) using MEME.

ChIP-seq in i-WT-r cells after 12 hr induction of EED led to H3K27me3 deposition as well as the presence of EED and SUZ12 at precisely those sites that retained H3K27me3 and PRC2 in the steady-state cage-mutant setting (Figure 2b, compare i-WT-r 12 hr with steady-state cage-MT; Figure 2c, compare lanes 1 and 8; Figure S4, compare panels e vs. f and h vs. i). We termed these “nucleation sites”, as they were the first regions where PRC2 was recruited and H3K27me3 deposited. Nucleation sites were grouped as either strong or weak with respect to the levels of PRC2 and H3K27me2/3 deposition at 12 hr of EED expression (left graphs in Figure 2d, e, f). Of note, these sites were specifically enriched for dense CpG islands (CGIs) (Figure 2c, lane 12). By 36 hr of EED induction in i-WT-r cells, H3K27me3 appeared at genomic regions distal from the initial nucleation sites, both at neighboring regions in cis and at distant intra-chromosomal regions, such that the steady-state WT distribution of H3K27me3 was approximated (Figure 2b, compare i-WT-r 36 hr with steady-state WT; Figure 2c, compare lane 3 with lane 10). These temporally established downstream sites of H3K27me3 deposition that emanated from an initiation site were termed “spreading sites”. Spreading sites were grouped as either proximal (-/+10 Kb) to at least one nucleation site (strong or weak) or distal and lacking a nucleation site in close proximity (Figure 2c, lane 1). In both cases, PRC2 levels were low as evidenced by the SUZ12 binding profile (Figure 2f).

WT EED rescue as a function of time revealed the dynamic build-up of H3K27me2 and -me3 domains. At 12 hr, H3K27me3 was deposited at the nucleation sites and mostly absent from the spreading sites (Figure 2b, i-WT-r 12 hr; Figure 2c, lane 1 and Figure 2d, left graph). H3K27me2 was also predominantly deposited at the nucleation sites, however; its presence was also detectable at spreading sites (Figure 2b, i-WT-r 12 hr; Figure 2c, lane 2 and Figure 2e, left graph). At 36 hr, H3K27me3 was detected at both nucleation and spreading sites, although the former were more enriched for this modification (Figure 2b, i-WT-r 36 hr; Figure 2c, lane 3 and Figure 2d, right graph). In contrast, H3K27me2 was depleted at nucleation sites and enriched at spreading sites (Figure 2b, i-WT-r 36 hr; Figure 2c, lane 4 and Figure 2e, right graph). As H3K27me3 levels accumulated over time in the i-WT-r, the H3K27me2 levels decreased at these same loci, displaying an anti-correlation between the two marks. This phenotype was similar to the steady-state WT scenario (Figure 2b, compare i-WT-r 36 hr with steady-state WT; Figure 2c, compare lanes 3, 4 with lanes 10, 11, respectively). Thus, deposition of H3K27me2 preceded that of H3K27me3 at the nucleation sites such that H3K27me2 was converted to H3K27me3 at 36 hr. Importantly and consistent with previous studies, H3K27me2 appropriately decorated intergenic regions and gene bodies (Ferrari et al., 2014; Lee et al., 2015), and not CGIs (Figure 2c, compare lanes 4 and 12; Figure S5a), while 71% of H3K27me3 spread +/-5 kb outside of the TSS.

The cage-mutant EED displayed very different dynamics in the rescue experiment. H3K27me3 deposition was severely compromised in the i-MT-r cells (Fig 2c, lane 6) and instead, H3K27me2 dynamics mirrored those of H3K27me3 in the WT rescue. At 12 hr, H3K27me2 was found at nucleation sites and was absent from spreading sites (Figure 2c, lane 5), and by 36 hr, H3K27me2 deposition intensified at nucleation sites, and inefficiently deposited at spreading sites as compared to WT EED rescue (Figure 2c, compare lane 4 with lane 7). Relative to the WT rescue, EED and SUZ12 were recruited at similar levels to nucleation sites in the cage-mutant rescue at 12 hr, but at reduced levels at 36 hr (Figure S4c-j and Figure 2f for SUZ12). In both cases, EED/SUZ12 occupancy at spreading sites in i-MT-r cells was minimal.

Taken together, our data suggest that the EED aromatic cage is not required for initial PRC2 recruitment and di-methylation, but is required for the efficient catalysis of the di-to tri-methyl reaction at nucleation sites. This outcome likely reflects both the increased stability of the WT EED on chromatin, imparted via the binding affinity of the aromatic cage for both di-and tri-methylated nucleosomes, and the allosteric activation of PRC2 via its binding to the tri-methylated version (Margueron et al., 2009). Defects in such binding affinity and in allosteric activation as in the i-MT-r case, result in H3K27me2 deposition at nucleation sites with subsequent, suboptimal H3K27me2 spreading, and the absence of a tri-methylated domain (Figure 2d and e). Notably, this profile contrasts in part with the steady-state case in which significant H3K27me3 deposition persisted at nucleation sites even without an intact aromatic cage (Figure 2c, compare lane 6 with lane 8). This discrepancy is not surprising as in contrast to the i-MT-r case that was devoid of pre-existing H3K27me3, there were pre-existing H3K27me3 nucleosomes at nucleation sites prior to the introduction of the cage mutation in the steady-state case. Therefore, PRC2 is more challenged in attaining appropriate *de novo* levels of H3K27me3 than in preserving those already established in the EED cage-mutant cells. Hence, at sites of active PRC2 recruitment (nucleation sites), the aromatic cage is required for *de novo* H3K27me3 domain formation, rather than its maintenance.

A motif analysis at strong nucleation sites revealed an overrepresentation of ‘GA’-rich and ‘GCN’ tandem repeat motifs (Figure 2g) relative to all CGIs in the genome. In total, 78% of strong nucleation sites had at least one of these motifs (Figure S5b). Furthermore, these motifs were unique relative to CGIs near the TSS of genes subject to spreading (Figure S5c). Although less significantly enriched, we did observe that similar ‘GA’-rich and ‘GCN’ tandem repeat motifs were overrepresented at the weak nucleation sites. However, the distribution of ‘GA’ content was slightly different and ‘GCN’ tandem repeats were shorter (Figure S5d).

To substantiate that PRC2 can indeed traverse the distance between nucleosomes to propagate its activity in cis, we reconstituted the spread of H3K27me3 from a neighboring pre-modified nucleosome *in vitro*. Oligonucleosome arrays were designed to contain a 2:6 ratio of effector:substrate histone octamer such that effector octamers contained either unmodified (-me0), pre-modified H3K27me3, or control K27A histones (Figure S6a). Substrate octamers were assembled using FLAG-H3 such that PRC2 histone methyltransferase activity on effector versus substrate nucleosomes could be distinguished. Increasing levels of PRC2 resulted in a stimulation of activity towards the substrate with pre-modified H3K27me3 (Figure S6b and c). Importantly, this stimulation was dependent on the aromatic cage of EED, as recombinant PRC2 harboring the EED Y365A mutation was ineffectual in this regard. Thus, the EED-H3K27me3 binding module underlies the spread of PRC2 activity in cis from an initial nucleation site.

### MTF2 along with JARID2 are necessary for *de novo* PRC2 targeting to chromatin

To identify factor(s) involved in the recruitment and stabilization of PRC2 on chromatin, we analyzed proteins enriched at the nucleation sites using an unbiased approach. We took advantage of the EED cage-mutant that failed to efficiently spread H3K27me2/3 and was stuck at the nucleation sites (Figure 2). Native ChIP-mass spectrometry (ChIP-MS) identified proteins associated with WT EED chromatin vs. EED cage-mutant chromatin (Figure 3a). In addition to the expected, core PRC2 subunits, proteins previously identified as PRC2 partners were enriched at the EED cage-mutant-containing nucleation sites (Figure 3b, Table S2). Amongst the top candidates, we identified JARID2, AEBP2, and MTF2, which were reported to have functional DNA binding activity, as well as roles in PRC2 recruitment and/or activity (Kim et al., 2009; Landeira et al., 2010; Li et al., 2010; Li et al., 2017; Pasini et al., 2010; Peng et al., 2009; Shen et al., 2009). Of note, elimination of these factors individually in steady-state mESCs did not completely disrupt PRC2 recruitment and had minimal effect on H3K27me3 levels. Importantly, MTF2 and JARID2 were shown to localize to GC-rich motifs (Li et al., 2017; Peng et al., 2009), which are similar to the GCN tandem repeat motif identified here as being enriched at the nucleation sites. We next gauged PRC2 recruitment and stability at nucleation sites in i-WT-r cells as a function of time of WT EED expression and the presence of JARID2/AEBP2/MTF2.

**Figure 3:**
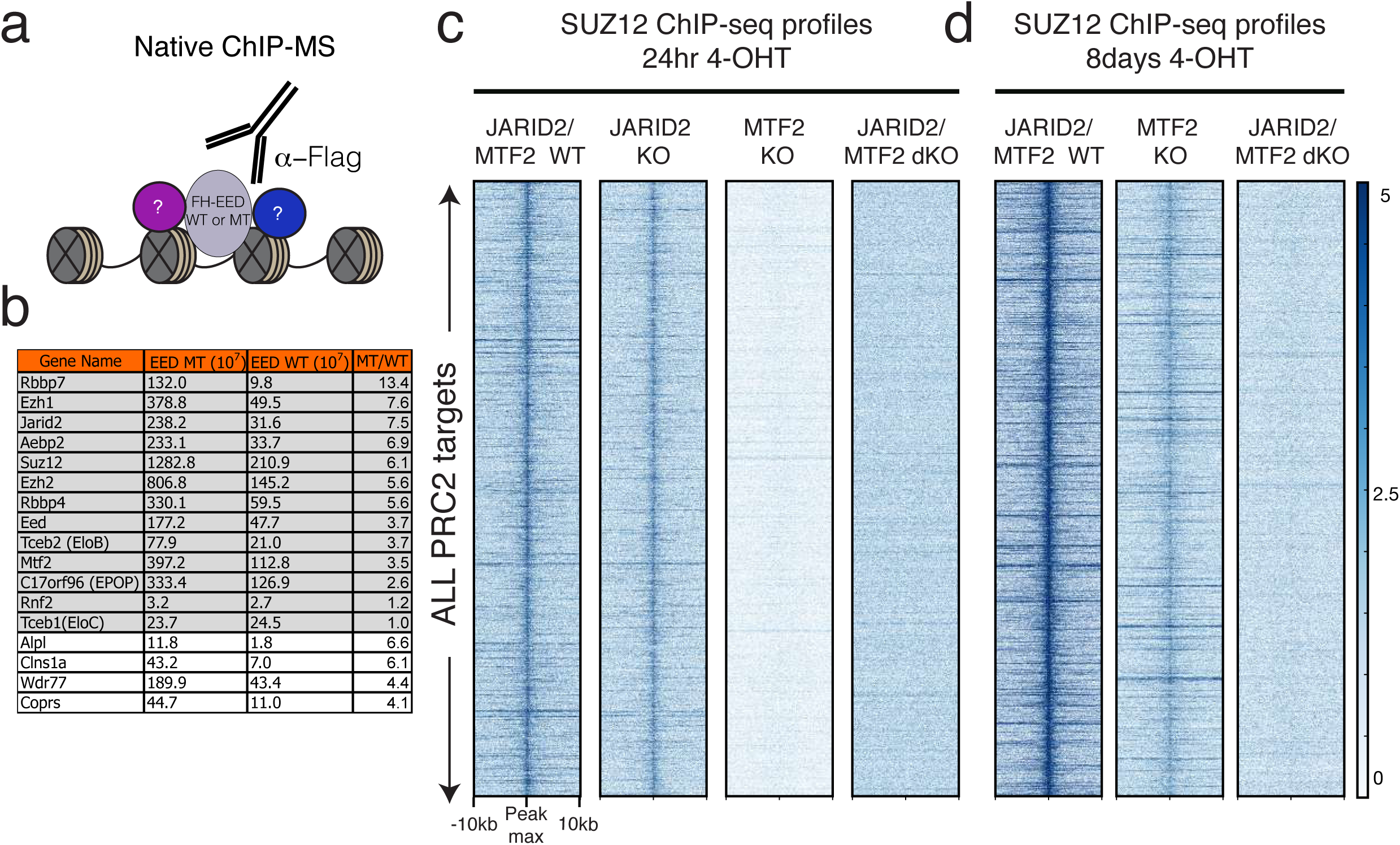
Deletion of MTF2 along with JARID2 abolishes *de novo* PRC2 targeting to chromatin. **a,** Scheme for native ChIP-MS approach to identify proteins enriched at chromatin comprising FLAG-HA tagged EED (FH-EED), either cage-mutant (Y365A) or WT. **b,** List of proteins enriched at the chromatin of EED cage-mutant vs. WT EED. To calculate the enrichment of specific proteins on EED cage-mutant bound chromatin vs. EED WT bound chromatin, peak integration values of peptides corresponding to identified proteins, were normalized to histones in each sample. The normalized values for individual proteins from the EED cage-mutant chromatin were divided by those from WT chromatin (MT/WT). Proteins highlighted in gray are core PRC2 components and previously reported PRC2 interactors. The last 4 proteins that are not highlighted have not been reported to interact with PRC2. **c, d,** Heatmaps of SUZ12, 24 hr **(c)** and 8 days **(d)** after 4-OHT treatment to rescue with WT EED, within a 20 kb window centered on the maximum value of peak signal in WT mESCs, comparing the genotypes indicated. (Scale: 0-5 for SUZ12).

When both JARID2 and MTF2 were eliminated in the i-WT-r cells, we observed a complete loss in the initial recruitment of PRC2 to nucleation sites (Figure 3c and d; Figure S5g and h). Consistent with our ChIP-MS data, ChIP-seq revealed that JARID2 and MTF2 were enriched at nucleation rather than spreading sites (Figure S5e and f). Elimination of JARID2 alone showed a mild effect on PRC2 recruitment, while elimination of MTF2 alone resulted in a loss of PRC2 recruitment to chromatin at 12 hr and 24 hr after EED expression (Figure S5h and h; Figure 3c). However, when the cells were cultured for longer periods and reached steady-state levels (8 days), we observed some recruitment of PRC2 in MTF2 KO cells, yet PRC2 recruitment was still undetectable in JARID2/MTF2 dKO cells (Figure 3d). Depletion of AEBP2 did not alter the *de novo* deposition of PRC2 activity in i-WT-r cells (data not shown). These results indicate that both JARID2 and MTF2 are necessary for the initial recruitment and stable binding of PRC2 to nucleation sites (see below). Of note, previous reports identified two major classes of PRC2 complexes (Grijzenhout et al., 2016), one comprising Jarid2/AEBP2 and the other comprising MTF2. Our results indicate that PRC2 candidates from either class can be recruited to nucleation sites through their JARID2 or MTF2 constituents.

### Deletion of a nucleation site delays H3K27me3 spreading

We postulated that nucleation sites might function to initiate the spread of PRC2 activity both locally and to regions that are genomically distant, but in close proximity as a result of 3D contacts. A purely cis-based propagation model would predict that cells lacking an intact nucleation site would no longer exhibit efficient deposition and spreading of H3K27me3 to neighboring regions. Thus, we generated an 8 kb genetic deletion in WT mESCs of a nucleation site identified on the *Evx2* promoter from which PRC2 activity spreads in cis across the *HoxD* cluster. The levels of H3K27me3 across the cluster remained unaffected by loss of the nucleation site (Figure S7a). However, pre-existing H3K27me3 adjacent to the deletion site might have been sufficient to recruit and stabilize PRC2 activity across the *HoxD* cluster. To control for this possibility, we generated the equivalent deletion of the *Evx2* nucleation site in uninduced i-WT-r cells, devoid of any adjacent H3K27me3 (Figure 4a). As expected, induced EED expression led to H3K27me3 deposition across the *HoxD* cluster, but importantly, in a delayed manner relative to cells possessing an intact nucleation site (Figure 4b). Deletion of two other nucleation sites resulted in similar delayed deposition and spreading of H3K27me3 to neighboring regions (Figure S7b and c). In each case, H3K27me3 levels were reduced relative to cells with the intact nucleation site after 24 hr of EED expression, but almost fully recovered by 36 hr (Figure 4b and Figure S7b, c). This delayed deposition was specific to regions in cis with respect to the deletion site, as H3K27me3 levels were restored to WT levels within just 24 hr at other intact nucleation sites (Figure S7d and e). Thus, nucleation sites can facilitate the spread of PRC2 activity in cis, but there exists a redundant network of interactions between genomic sites, pointing to a mechanism by which only a small number of nucleation sites can lead to the formation of thousands of H3K27me2/3 domains.

**Figure 4:**
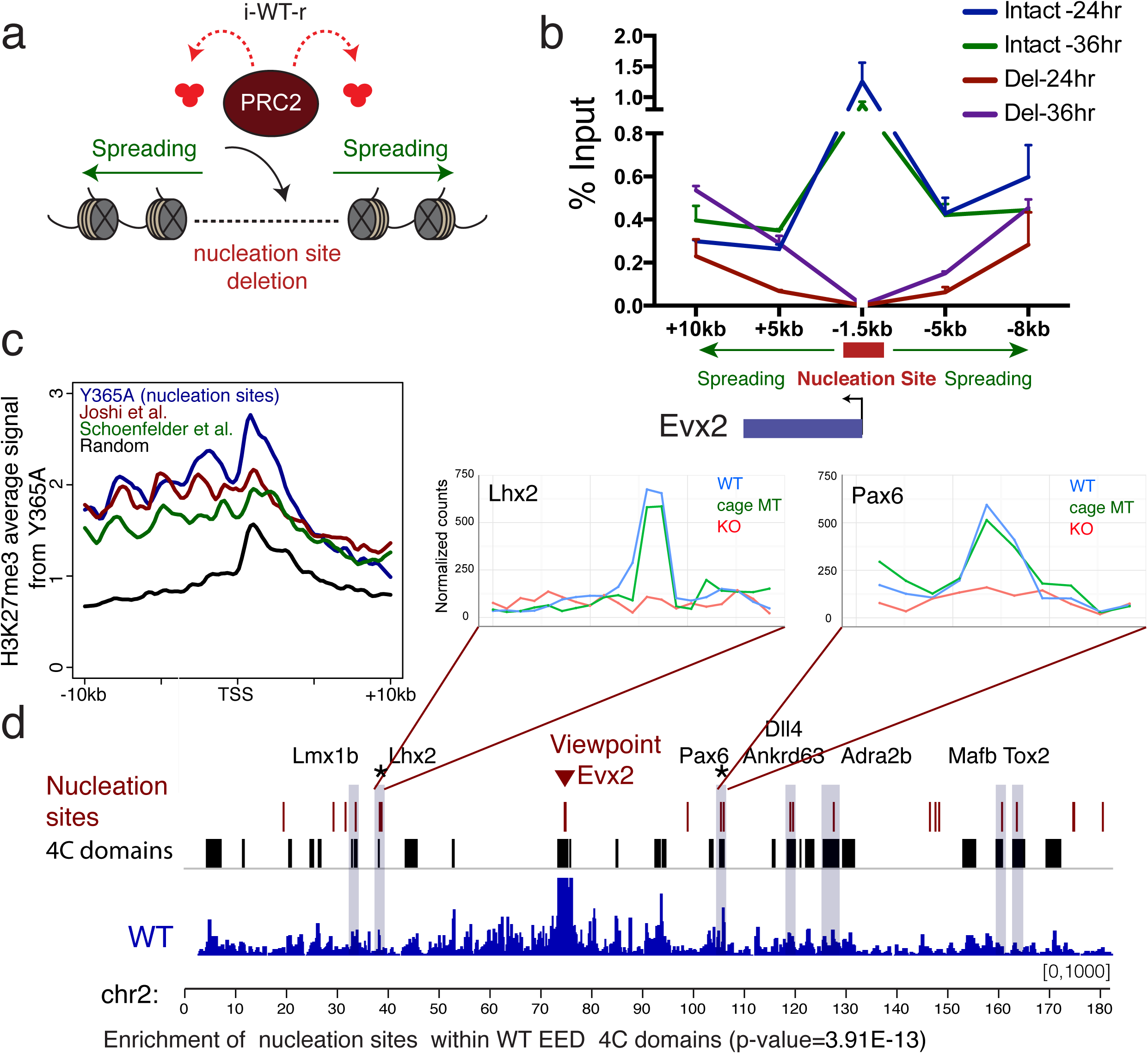
Functional and structural characterization of nucleation sites. **a,** Approach to assay for the formation of H3K27me3 domains in the absence of a nucleation site. The *Evx2* nucleation site (red) is deleted within the i-WT-r rescue system (Figure 2a), and spreading (green) of PRC2 activity in cis was assayed by ChIP-qPCR. **b,** H3K27me3 ChIP-qPCR in i-WT-r cells harboring an intact versus deleted nucleation site, at the positions indicated relative to the TSS of *Evx2* (n=2). WT EED rescue was performed with 4-OHT treatment for 24 hr and 36 hr, as indicated. Enrichment is calculated as % of input. **c,** Average H3K27me3 ChIP-seq density in cage-mutant EED (Y365A) mESCs was plotted within a 20 kb window centered around the TSS of the top 293 H3K27me3 target genes (closest genes to 200 peaks identified in Figure 1d) in these cells (blue, positive control). Similarly, H3K27me3 density was plotted across a set of spatially interacting polycomb target genes from Joshi *et al*. (red, 186 genes), Schoenfelder *et al*. (green, 195 genes), and across a random set of H3K27me3 target genes (black, 359 genes), in WT mESCs. **d,** 4C-seq track depicting the interaction frequency with respect to the *Evx2* (strong nucleation site) viewpoint. 4C domains (in black) and nucleation sites (in red) are indicated above. 4C domains represent the intersection of 2 replicate experiments. Overlap between nucleation sites and 4C domains are highlighted in gray. Nucleation sites are significantly enriched within 4C domains interacting with *Evx2* (p-value=3.91E-13). Nucleation sites having reduced interactions with the *Evx2* viewpoint in the EED KO but not cage-mutant (Y365A) or WT setting, are labeled with an asterisk and their interaction profiles are plotted (top right).

### Nucleation sites spatially interact in the nucleus

3C-based approaches have recently shown that polycomb targets form a cluster of highly enriched H3K27me3 regions in mESCs (Denholtz et al., 2013; Joshi et al., 2015; Schoenfelder et al., 2015; Vieux-Rochas et al., 2015) as a result of long-range interactions. As this chromatin organization could conceivably orchestrate the long-distance spread of PRC2 activity, we next explored whether the nucleation sites we identified were enriched within published sets of spatially interacting polycomb targets. We first plotted H3K27me3 density in the Y365A cage-mutant cells, centered around the TSS of genes reported by two independent groups to lie in close spatial proximity. Strikingly, the average H3K27me3 density across these genes was similar to the positive control set of nucleation sites (Y365A), and was enriched over a randomized negative control (Figure 4c). Furthermore, another group recently identified 10 genomic sites via 4C-seq that frequently interact with the *HoxD13* locus (Vieux-Rochas et al., 2015), which lies adjacent to the *Evx2* nucleation site. Of these reported *HoxD13* interactions, 70% overlapped with the set of nucleation sites we identified (data not shown). Thus, nucleation sites likely represent a large portion of the polycomb target interaction network that is formed in mESCs.

To confirm the spatial proximity of nucleation sites, we performed 4C-seq in WT mESCs using nucleation sites as viewpoints. In agreement with our model, the *Evx2* nucleation site was significantly enriched for interactions with other intra-chromosomal nucleation sites in WT mESCs (Figure 4d, p-value=3.91E-13). We confirmed the spatial proximity of nucleation sites with 4C-seq using four other nucleation sites as viewpoints: *Lhx2, Cyp26b1, HoxA*, and *HoxB* clusters (Table S3). As controls, we used viewpoints placed at each of three different spreading sites (*HoxC* cluster, *Specc1*, and *Irak4*), the active gene *Pou5f1*, and a gene desert, which did not demonstrate significant interactions with PRC2 nucleation sites on the same chromosome (Table S3). These observations suggest that spatial clustering can facilitate the spreading of H3K27me3 activity via long-range chromatin contacts such that one nucleation site can compensate for the absence of another, albeit with delayed dynamics (Figure 4b and Figure S7b, c).

We next asked whether chromatin contacts of nucleation sites are dependent on polycomb activity itself. We observed no significant changes in the 4C interactome in WT versus cage-mutant EED (Y365A) cells across all 11 nucleation sites that interacted with *Evx2*. However, 2 nucleation sites (*Lhx2* and *Pax6*) had a significantly reduced interaction frequency in EED KO cells as compared to WT mESCs (Figure 4d, top right). Taken together, our data strongly suggest that nucleation sites are in spatial proximity to each other, and that consistent with recent reports, this sub-nuclear architecture is partially dependent on polycomb repressive chromatin domains (Denholtz et al., 2013; Schoenfelder et al., 2015).

### PRC2 spreading from a group of foci to proximal and distal regions on chromatin

Our 4C data demonstrating the spatial proximity of PRC2 nucleation sites can be seen as a molecular representation of polycomb bodies within the nucleus. To visualize the progression of H3K27me3 foci formation, we performed immunofluorescence in i-WT-r cells using antibodies against H3K27me3 and EZH2, at 12 hr increments after EED expression was induced. Indeed, we were able to detect H3K27me3 foci starting just 12 hr after EED induction (Figure 5a), which corresponds to the initial appearance of H3K27me3 peaks by ChIP-seq (Figure 2b, c). ImmunoFISH experiments confirmed that these 12 hr foci overlapped with a nucleation site (*Evx2,* near *HoxD*) at a significantly higher frequency than with a spreading site (*HoxC*) (Figure 5b, Figure S8 and Table S4). Importantly, *Evx2* alleles overlapped with denser H3K27me3 foci, as compared to *HoxC* alleles (Figure S8 and Table S4). The observed foci (Figure 5a), increased in number and size by 24 hr, and eventually spread to large regions of the nucleus by 36 hr. EZH2 exhibited a diffuse staining pattern, likely due to its promiscuous binding to nascent RNA transcripts (Davidovich et al., 2013; Kaneko et al., 2013) and/or its dynamic interaction with these foci, as evidenced by *Drosophila* FRAP experiments in which Polycomb group (PcG) proteins exhibited short residency times on chromatin in stem cells (Fonseca et al., 2012). Thus, the earliest phase of PRC2 recruitment occurred at a limited discrete group of foci, consistent with our model of nucleation followed by spreading to proximal and distal regions.

**Figure 5:**
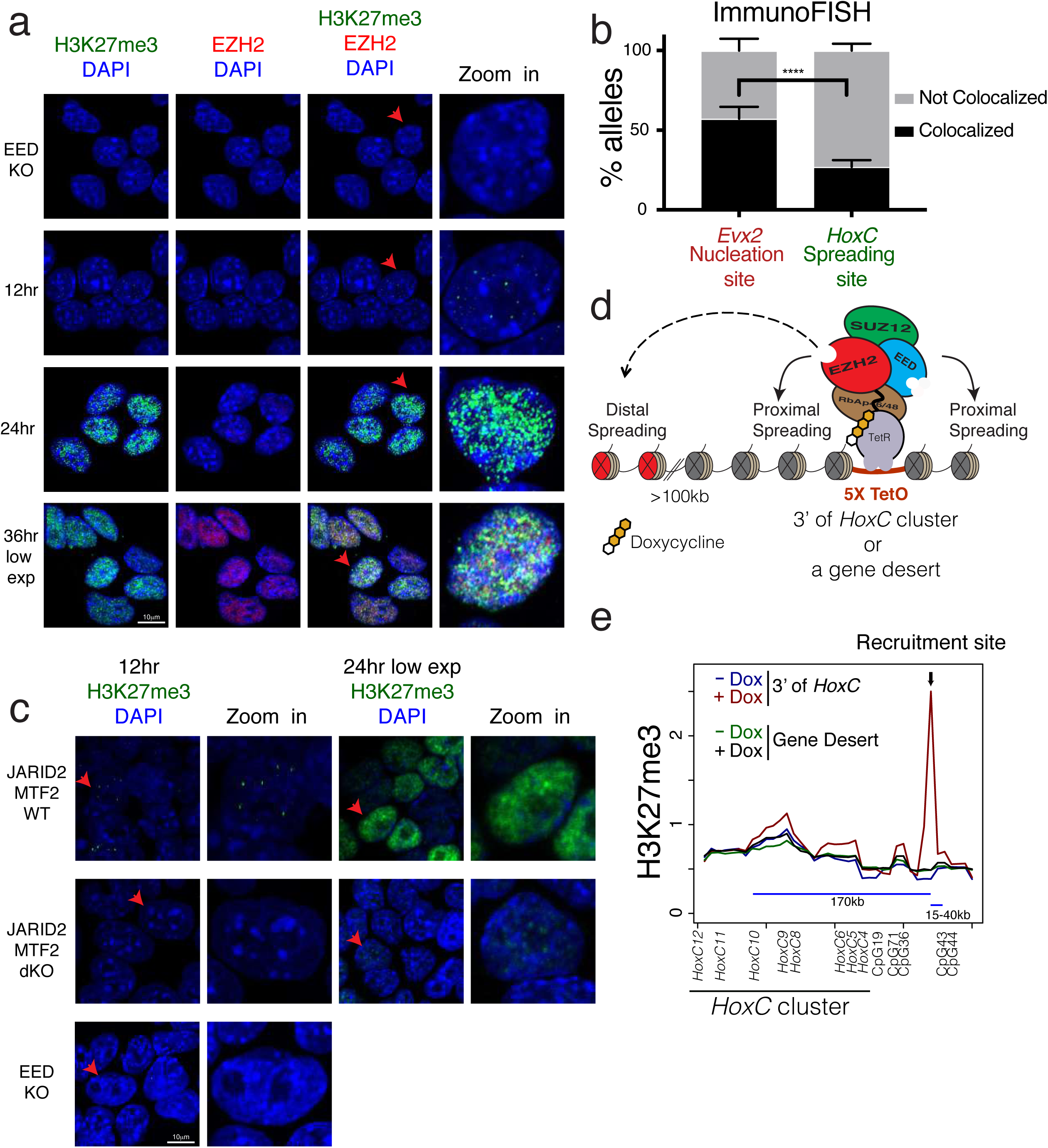
H3K27me3 domains initiate within nuclear hubs of PRC2 activity, from which H3K27me3 spreads in cis and far-cis via long-range interactions. **a,** Immunofluorescence using the indicated antibodies at 0, 12, 24 and 36 hr after rescue of WT EED expression in i-WT-r mESCs. H3K27me3 staining at 36 hr is shown at lower exposures. The rightmost panel is a zoomed-in image of the cell labeled with a red arrow. **b,** Quantification of % alleles colocalized with H3K27me3 foci using ImmunoFISH, comparing probes for a nucleation site (*Evx2*) and a spreading site (*HoxC*) at 12 hr after WT EED expression in i-WT-r cells. n for *Evx2*: 98, n for *HoxC*: 100. Fisher‘s exact test of a combination of two biological replicates: ****p-value<0.0001. **c,** Immunofluorescence using H3K27me3 antibodies at 12 hr (left) and 24 hr (right) after rescue of WT EED expression in i-WT-r mESCs with the indicated genotypes. H3K27me3 staining at 24 hr is shown at lower exposures. The rightmost panel is a zoomed-in image of the cell labeled with a red arrow. Staining of EED KO cells (0 hr, without 4-OHT) is shown at bottom left as a control. **d,** Scheme for ectopic targeting of PRC2 using the Tet repressor and Tet operator system in i-WT-r cells (see text for details). **e,** Average ChIP-seq read density of H3K27me3 plotted around the recruitment site using 20 kb bins before (-Dox) and after (+Dox) the induction of PRC2 recruitment downstream of the *HoxC* cluster and 4-OHT treatment for 24 hr to induce expression of WT EED. *HoxC10* spatially interacted with the recruitment site downstream of the *HoxC* cluster and with the genes indicated and with CpGs, as determined by 4C-seq.

We performed similar analyses using i-WT-r cells having a KO of both JARID2 and MTF2 and rescued with WT EED expression. As expected, H3K27me3 foci formation in JARID2/MTF2 dKO cells were undetectable after 12 hr rescue (Figure 5c). Importantly, delayed H3K27me3 foci formation was detected in JARID2/MTF2 dKO cells after 24 hr of rescue. However, we failed to detect (stable) PRC2 recruitment to chromatin in JARID2/MTF2 dKO cells, even after 8 days of rescue with WT EED (Figure 3c, d). This result strongly suggests that PRC2 can be transiently recruited to nucleation sites (a “hit & run” mechanism, see below) and thereby generate H3K27me3-forming foci, yet fail to be stabilized on chromatin in the absence of JARID2 and MTF2.

To examine the distal spreading from initial nucleation sites within H3K27me3 foci, we devised an artificial system in which PRC2 can be recruited to a region with known spatial interactions lacking H3K27me3. Using CRISPR/Cas9 in i-WT-r cells, a cassette containing 5 copies of the Tet operator sequence (5xTetO) was integrated within a gene dense region that interacts broadly across the *HoxC* cluster, sitting ∼150 kb downstream (located at chromosome 15). As a control, the same cassette was inserted within a gene desert on chromosome 5 that had no detectable interactions with the *HoxC* cluster (Figure 5d). Both endogenous *EZH2* alleles were tagged with a mutant version of the Tet-repressor (TetR) DNA binding domain that binds to inserted TetO sequence arrays such that PRC2 recruitment could be induced (Figure 5d). After doxycycline addition, expression of WT EED at 24 hr resulted in the formation of H3K27me3 domains of ∼5 kb in size at both artificial recruitment sites in i-WT-r cells (Figure S9). Importantly, recruitment of TetR-EZH2 downstream of *HoxC*, which is in contact with *HoxC*, but not to the gene desert, led to a specific and significant increase in H3K27me3 signal on the entire *HoxC* locus (Figure 5e). These observations support our model that PRC2 activity spreads from an initial nucleation site not only to proximal regions in cis, but also to distal, contacting regions.

We next took advantage of the published data from Zheng et al., 2016 (Zheng et al., 2016), that mapped H3K27me3 domains during development. They showed that H3K27me3 was depleted from promoters in preimplantation embryos and *de novo* H3K27me3 was deposited in the epiblast, indicating a massive epigenetic reprogramming. We examined whether our nucleation sites overlapped with the *de novo* H3K27me3 peaks observed in the E5.5 epiblast. Indeed, the proportion of nucleation sites that significantly overlap with H3K27me3 peaks increased progressively during the transition from the 8-cell to the ICM, and finally to the E5.5 epiblast stage Figure S10a, b, and c). Strikingly, almost all the PRC2 nucleation sites overlap with those of the E5.5 epiblast. However, there were considerably more *de novo* peaks at the E5.5 epiblast stage that did not overlap with nucleation sites (Figure S10c), suggesting that spreading from the nucleation sites had already begun. Interestingly, polycomb deposition at nucleation sites occurs as early as the ICM stage (Figure S10d), suggesting that the critical chromatin rewiring decision is made at this phase. These observations provide evidence that the two-step mechanism that we detail below is applicable during *in vivo* epigenetic reprogramming.

## Discussion

Taken together, our findings support the following model in which polycomb foci are formed at a subset of polycomb target genes (Figure 6) that are predominantly developmentally regulated and maintained in a silent state in mESCs. Nucleation sites comprising specific CGIs in general can engage in the nucleus to initiate a network of interactions partially dependent on PRC2 and its product, H3K27me3. From these initial nucleation sites, PRC2 spreads H3K27me2/3 to neighboring regions proximally as well as distally via long-range 3D contacts, all within sub-nuclear foci of polycomb activity. PRC2 recruitment and stability on chromatin is dependent on its on- and off-rates, shown by arrows pointed “to” and “from” PRC2 recruitment site(s), respectively, in Figure 6. PRC2 alone may interact with the nucleation sites through a “hit & run” mechanism; an unstable interaction such that its off-rate is higher than its on-rate. Its on-rate is increased when associated with JARID2, and further increased when associated with MTF2. Once PRC2 reaches sufficient concentrations, it catalyzes H3K27me2 first and converts it to H3K27me3 at the nucleation sites, where it is more stably bound. From this initial nucleation event, PRC2 rapidly spreads H3K27me2/3 domains across the genome, such that its residency time on chromatin is decreased as it moves further from the nucleation sites, resulting in a more pronounced deposition of H3K27me2 than H3K27me3. The cage of EED is required for the stability and increased trimethylation activity of PRC2 through its interaction with H3K27me3 and thus, for efficient spreading of both H3K27 di-and trimethylation.

**Figure 6:**
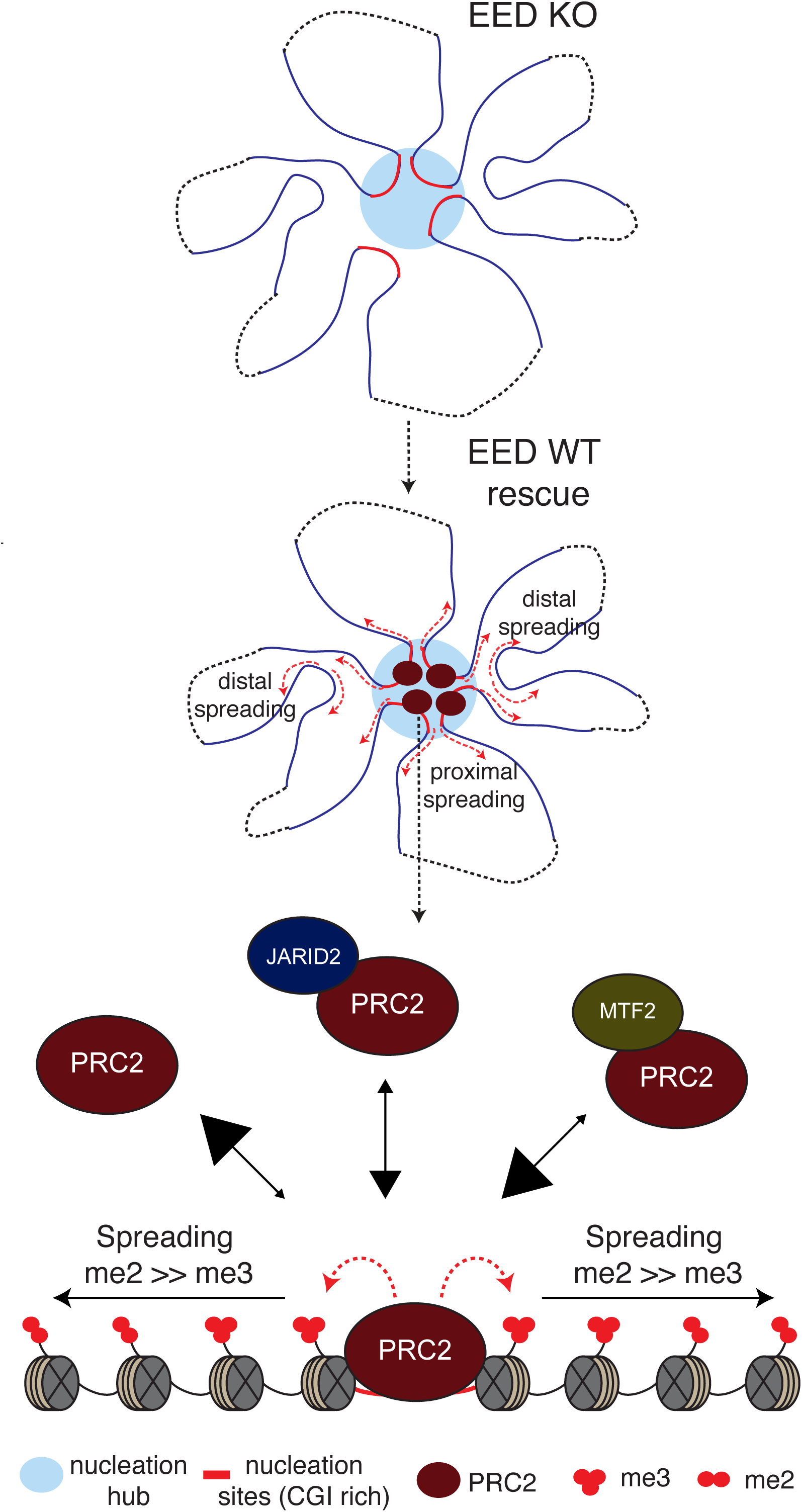
Model for PRC2 recruitment to chromatin and establishment of H3K27me2/3 domains. In EED KO mESCs, spatial interactions among some, but not all nucleation sites are lost. Upon expression of EED, PRC2 is first recruited to regions called “nucleation hubs” that have a specific subset of CGIs forming spatial clusters in the nucleus. The stable binding of PRC2 to chromatin depends on its on- and off-rates, shown by arrows pointed “to” and “from” chromatin, respectively, with the size of the arrowheads reflecting relative PRC2 binding. PRC2 alone exhibits the lowest stability on chromatin, but its stability is increased when complexed with Jarid2, and further increased when complexed with MTF2. Upon reaching sufficient concentrations, PRC2 catalyzes H3K27me2 first and converts it to H3K27me3 at the nucleation sites, where it is more stably bound. From this initial nucleation event, PRC2 rapidly spreads H3K27me2/3 domains across the genome, such that its residency time on chromatin is decreased as it moves further from the nucleation sites, resulting in a more pronounced deposition of H3K27me2 than H3K27me3 to neighboring regions proximally as well as distally via long-range 3D contacts. Interaction between the cage of EED and H3K27me3 is required for allosteric activation of PRC2 and for the stability of PRC2 on chromatin and thus, for efficient spreading of both di-and trimethylation of H3K27.

Attempts to identify an analog of the PRE-based recruitment paradigm from *Drosophila* (Cuddapah et al., 2012; Mendenhall et al., 2010; Schorderet et al., 2013; Sing et al., 2009; Woo et al., 2010) for the mammalian polycomb system have been challenging. Two putative mammalian PREs have been identified – a 1.4 kb region on mouse *HoxD10* (Schorderet et al., 2013) and a 1.8 kb region between human *HoxD11* and *HoxD12* (Woo et al., 2010). While both of these regions retained H3K27me3 in the cage-mutant setting, their levels were significantly lower than in WT cells, unlike the neighboring *Evx2* nucleation site. This finding suggests that *Evx2* may be the strong “PRE” from which PRC2 activity spreads in cis through the *HoxD* cluster. Though specific mammalian PREs have remained elusive, genome-wide studies have identified correlations between PRC2 targets and CGIs, raising the possibility that these sequences influence recruitment (Ku et al., 2008; Mohn et al., 2008). Indeed, exogenous insertion of GC-rich elements in mESCs was sufficient to recruit PRC2 activity (Lynch et al., 2012; Mendenhall et al., 2010). However, this effect requires that a given CGI is devoid of activating motifs (Mendenhall et al., 2010). In accordance, PRC2 binds only to CGIs of non-transcribed genes (Riising et al., 2014). Interestingly, despite its high density CGIs, the *HoxC* cluster did not retain H3K27me3 in the cage-mutants (Figure 1e), classifying it as a spreading site. Nonetheless, we have identified GCN motifs within CGIs near *HoxC9* and *HoxC12,* suggesting that these CGIs may serve as nucleation sites to spread PRC2 activity through the *HoxC* cluster. However, *HoxC12* was devoid of H3K27me3 and decorated by RNA Pol II, indicating that it is transcriptionally active. On the other hand, the CGI near *HoxC9* was decorated by an active mark, H3K27ac, flanked by H3K27me3, implying that it is prone to activation. These observations reinforce our previous findings that transcriptional activity and/or accompanying active chromatin features may counteract polycomb nucleation and spreading (Narendra et al., 2015).

Recent reports proposed that heterochromatin domain formation is driven by liquid phase-separation (for review see Hyman et al., 2014 (Hyman et al., 2014)), which is induced by a high local concentration of HP1α, in *Drosophila* and mammals (Larson et al., 2017; Strom et al., 2017). Such a mechanism may pertain to the formation of sub-nuclear foci comprising nucleation sites with high, local concentrations of PRC2/MTF2 and/or PRC2/Jarid2 (Figure 2f, Figure S5e and f). This assemblage could drive the effective catalysis of H3K27me2/3.

Our finding that JARID2 and MTF2 are crucial for full recruitment of PRC2 to chromatin and for the establishment of *de novo* chromatin domains, provides insights as to how metastable transcriptional states are established in normal and disease states in response to developmental and environmental signals. The approaches used herein may pioneer future studies aimed at identifying the mechanisms underlying the establishment of other features of Polycomb domains as well as other types of chromatin domains.

## Acknowledgements

We thank Drs. L. Vales, K. J. Armache and E. O. Mazzoni for comments on the manuscript, J. Granat for technical assistance, Drs. R.A. Ganai, T. Escobar, and S. Krishnan for discussions, Y. Grobler for providing *Drosophila* S2R+ cells, Dr. A. Sfeir for kind gift of the C57BL/6 mESCs, New York University Langone Medical Center (NYULMC) Genome Technology Center, particularly A. Heguy, P. Zappile, and P. Meyn, for help with sequencing, NYULMC Cytometry and Cell Sorting Core for help with FACS, and NYULMC Microscopy Core for help with imaging. The NYULMC Genome Technology Center and the NYUMC Cytometry and Cell Sorting Core are partially supported by the Cancer Center Support Grant, P30CA016087 at the Laura and Isaac Perlmutter Cancer Center. This work utilized computing resources at the High Performance Computing Facility of the Center for Health Informatics and Bioinformatics at the NYULMC. P.R.R. is a National Cancer Center and American Society of Hematology Fellow. This work was supported by grants from the National Cancer Institute (9R01CA199652-13A1 to D.R.), the Howard Hughes Medical Institute (to D.R.), the National Institutes of Health (R01GM086852 and R01GM112192, to J.A.S.; GM110174 and CA196539 to B.A.G.).

## Author Contributions

O.O., V.N., and D.R. conceived the project, designed the experiments, and wrote the paper; O.O., performed most of the experiments and the bioinformatic data analysis; V.N. helped with designing the initial CRISPR constructs, analyzed RNA-seq and the initial ChIP-seq data; G.L., K.K., and B.A.G. quantified histone modifications and helped with ChIP-MS; C.L. prepared the hetero-oligonucleosomes and performed the *in vitro* histone methyltransferase assay; N.D. helped with the bioinformatic data analysis; R.R. analyzed the 4C-seq data; P.R.R. and J.A.S. advised on 4C-seq procedure and analysis. L. B. performed the immunoFISH experiment.

The authors declare no competing financial interests. Correspondence and requests for materials should be addressed to D.R.

## Note

The design of the experiments described herein to detect PRC2 nucleation sites, domain formation and most of the results and conclusions were presented approximately two and a half years ago at a chromatin meeting. Solely to set the record.

## Supplemental Figure Legends

**Figure S1:**
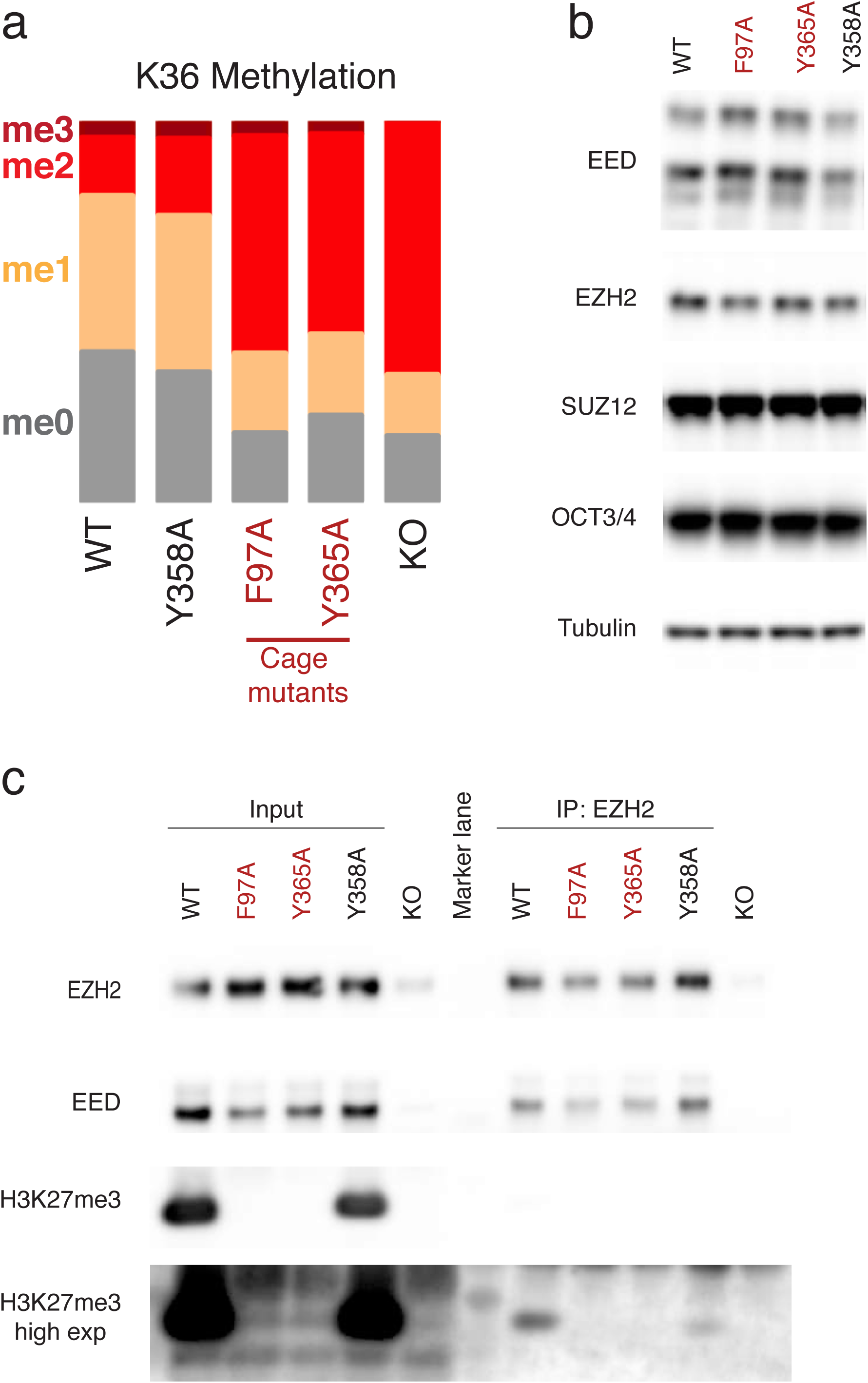
EED cage-mutant mESCs have elevated levels of H3K36me2 and intact PRC2 complex. **a,** Methylation status of H3K36 in mESCs of the indicated genotypes as identified by quantitative histone mass-spectrometry. **b,** Western blot using the indicated antibodies on whole cell extracts from mESCs. **c,** Immunoprecipitation (IP) of EZH2 using nuclear extracts from mESCs. Western blot was performed on input (left) and IP (right) samples with the antibodies indicated. Cage-mutants are highlighted in red.

**Figure S2:**
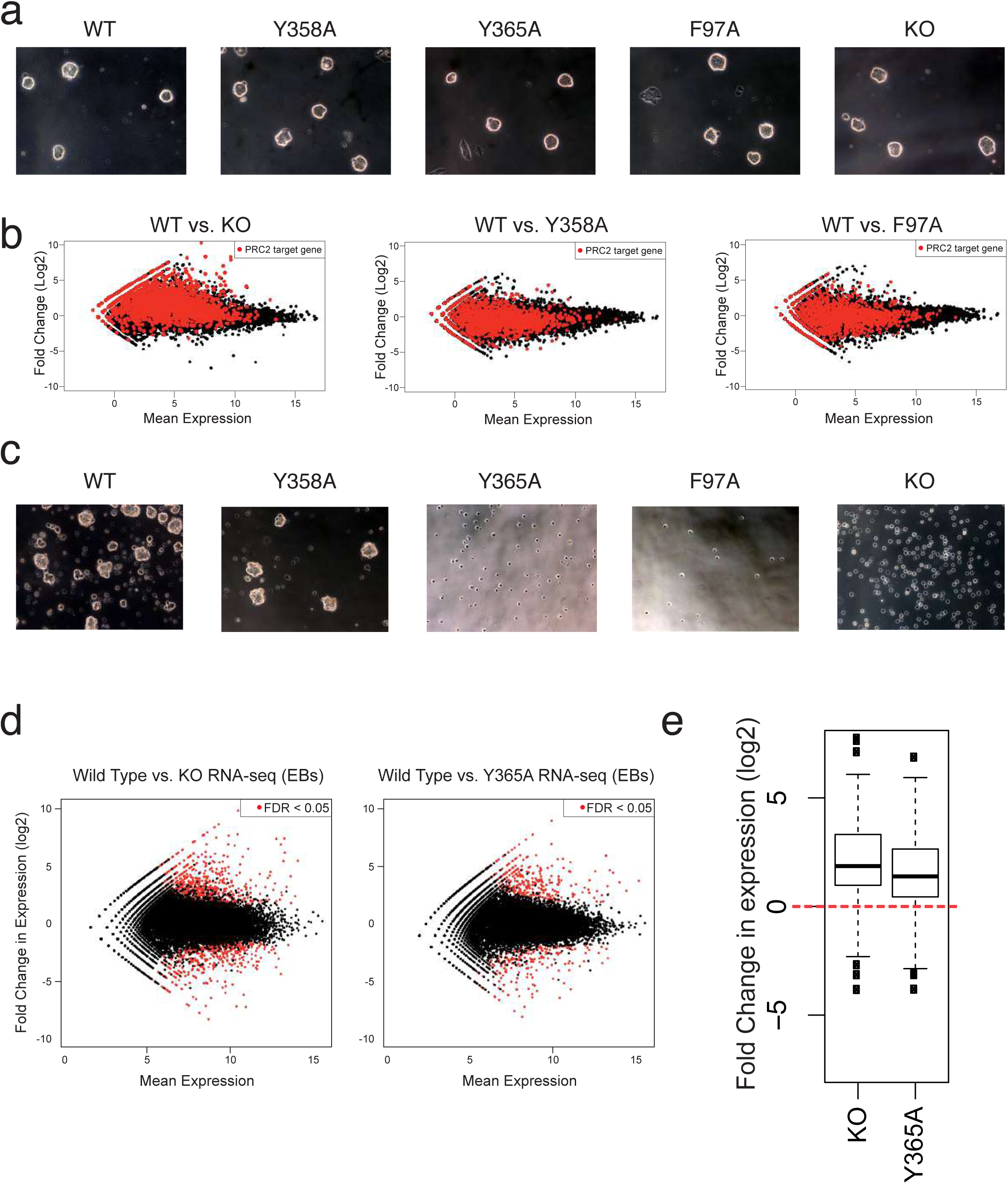
Cage-mutant EED mESCs maintained appropriate patterns of gene expression, but failed to differentiate. **a,** Bright-field microscopy images of undifferentiated mESCs comparing the indicated genotypes of EED. **b,** MA plots of RNA-seq data comparing mESCs of the indicated genotypes of EED. Each dot represents a gene. The x-axis represents its mean abundance across samples, and the y-axis represents the log2 fold enrichment between samples. PRC2 targets are colored in red. **c,** Bright-field microscopy images of mESCs differentiated into the cardiac lineage. Images were taken 3 days after differentiation. **d,** MA plots of RNA-seq data within 2 days of differentiation into embryoid bodies (EBs) of mESC with the genotypes of EED indicated. Differentially expressed genes (FDR<0.05) are colored in red. **e,** Box plot of genes that were repressed in a WT setting within 2 days of differentiation of mESCs into embryoid bodies (EBs) (*de novo* targets of PRC2). Y-value represents the relative expression of these genes in EED KO and Y365A EBs, relative to the WT background.

**Figure S3:**
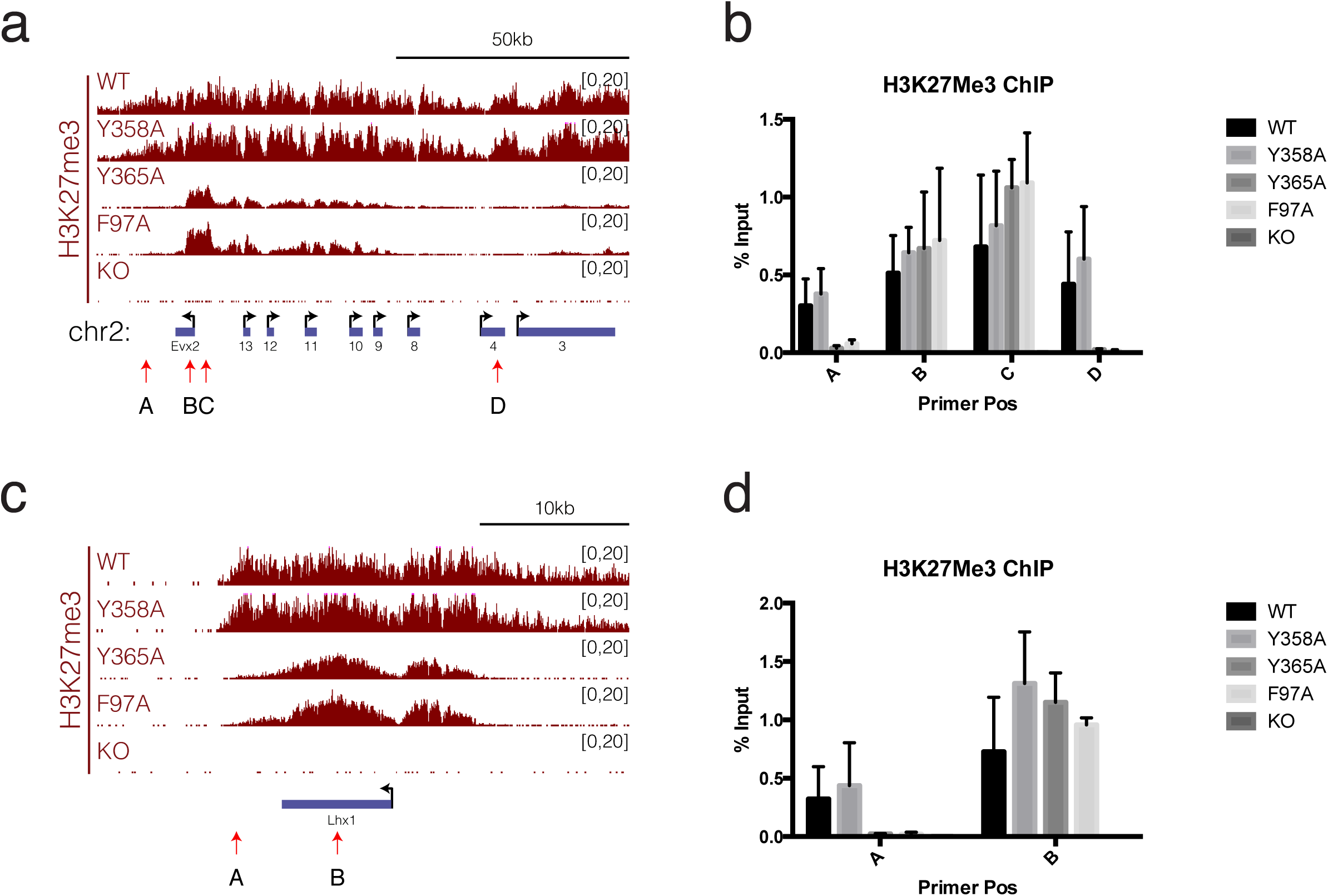
ChIP-qPCR validated H3K27me3 ChIP-seq results in the EED cage-mutants. **a, c** ChIP-seq tracks for H3K27me3 along the *Evx2* **(a)** and *Lhx1* **(c)** gene in mESCs with the indicated genotypes of EED. Positions of primers used in ChIP-qPCR are indicated by red arrows. **b, d,** H3K27me3 ChIP-qPCR using the primers from **a** and **c**, respectively. Enrichments were calculated as % of input. All ChIP-qPCR experiments were performed in two biological replicates.

**Figure S4:**
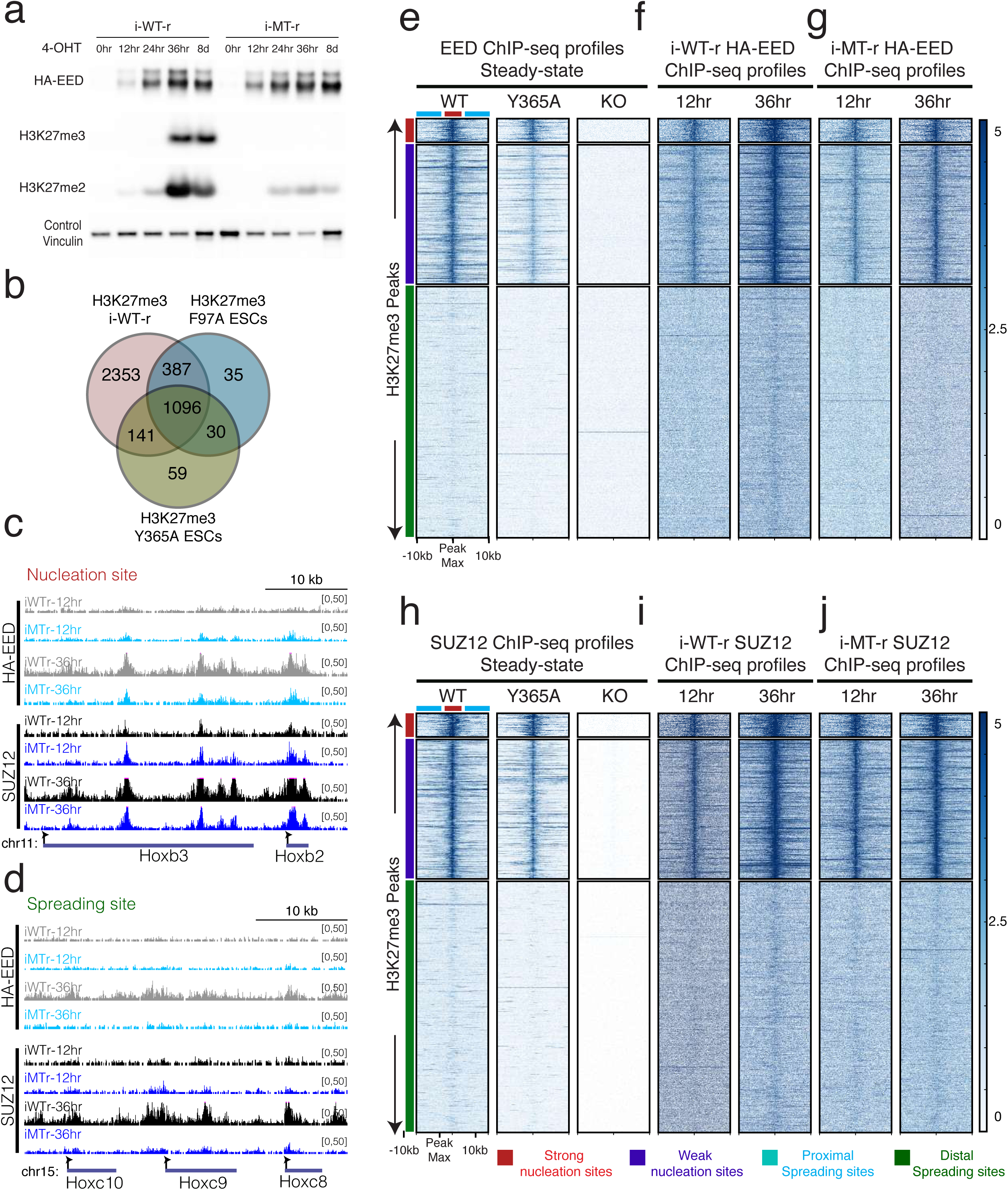
Initial deposition of H3K27me3 and PRC2 in i-WT-r and i-MT-r mESCs strongly overlapped with those sites that retained H3K27me3 and PRC2 in the steady-state EED cage-mutants. **a,** Validation of the i-WT-r and i-MT-r systems. Western blot using indicated antibodies on whole cell extracts after 4-OHT treatment to induce expression of EED, either WT (i-WT-r) or cage-mutant (i-MT-r), at the time points indicated. **b,** Venn diagram showing the overlap of H3K27me3 peaks between the 12 hr i-WT-r, Y365A and F97A mESCs. **c**, **d**, ChIP-seq tracks for HA-EED and SUZ12 along the *HoxB* cluster (nucleation sites) **(c)**, or the *HoxC* cluster (spreading sites) **(d)** in i-WT-r and i-MT-r cells after 4-OHT treatment at the time points indicated. **e**, Heatmaps of EED ChIP-seq performed on the indicated genotypes of EED in steady-state. **f, g**, Heatmaps of HA-EED ChIP-seq upon EED rescue, either WT **(f)** or cage-mutant **(g)** by 4-OHT treatment for the specified time points. **h**, **i**, **j**, Heatmaps of SUZ12 ChIP-seq performed on the indicated genotypes of EED in steady-state **(h)** or upon rescue with EED, either WT **(i)** or cage-mutant **(j)** with 4-OHT at the specified time points, within a 20 kb window centered on the maximum value of peak signal in WT mESCs. H3K27me3 peaks were sorted in descending order by signal intensity at the 12 hr time-point. Strong nucleation sites are marked in red (total=237), weak nucleation sites in blue (total=1389), proximal spreading sites in cyan and distal spreading sites in green (total= 36618, representative of randomly selected 2500 peaks are used for the Heatmaps). (Scales: 0-5 for EED, HA-EED and SUZ12).

**Figure S5:**
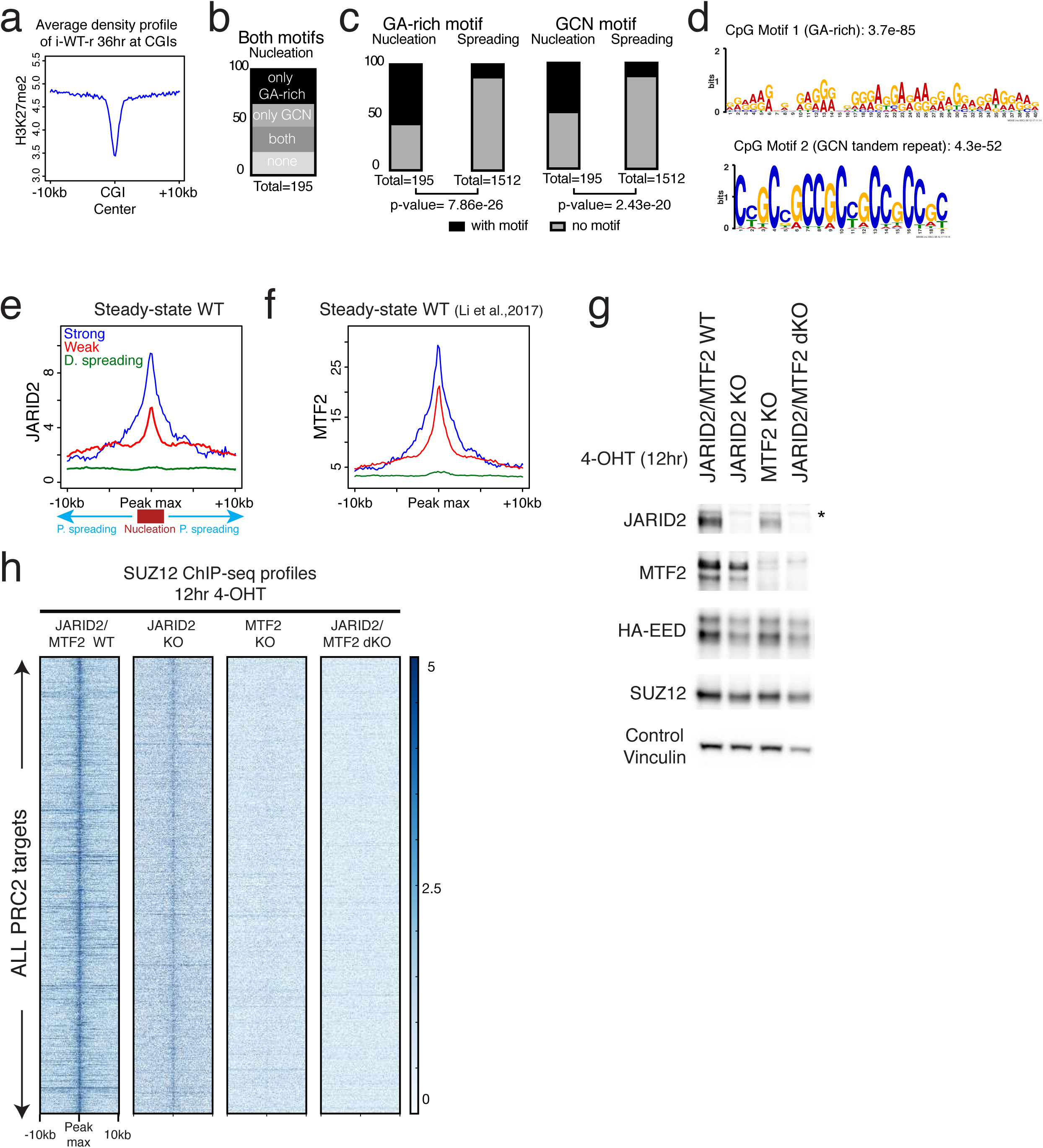
Features associated with nucleation versus spreading site CGIs. **a,** Average H3K27me2 density profile of the i-WT-r at 36 hr within CGIs. **b**, % of nucleation sites possessing either or both of the GA-rich and GCN motifs. **c**, % of nucleation and spreading sites containing either the GA-rich (left) or GCN tandem repeat motif (right). P-values represent the enrichment of these motifs within CGIs of nucleation versus spreading sites. **d**, Motif analysis of randomly selected 200 weak nucleation site CGIs (total=1085) using MEME. Iteration of the random selection resulted in identification of similar motifs. **e, f,** Average density profiles of JARID2 **(e)** or MTF2 (from Li *et al.,* 2017) **(f)** on nucleation sites (strong and weak) and spreading sites (D=distal, P=proximal), within a 20 kb window centered on maximum value of peak signal in WT mESCs. **g**, Western blot using the indicated antibodies on whole cell extracts derived from i-WT-r cells with the indicated genotypes after 12 hr of 4-OHT treatment. Bands exhibiting different sizes in the case of MTF2 and of HA-EED represent different isoforms of the proteins. Non-specific bands are labeled with an asterisk. **h**, Heatmaps of SUZ12 after 12 hr of 4-OHT treatment, within a 20 kb window centered on the maximum value of peak signal in WT mESCs comparing the indicated genotypes. (Scale: 0-5 for SUZ12).

**Figure S6:**
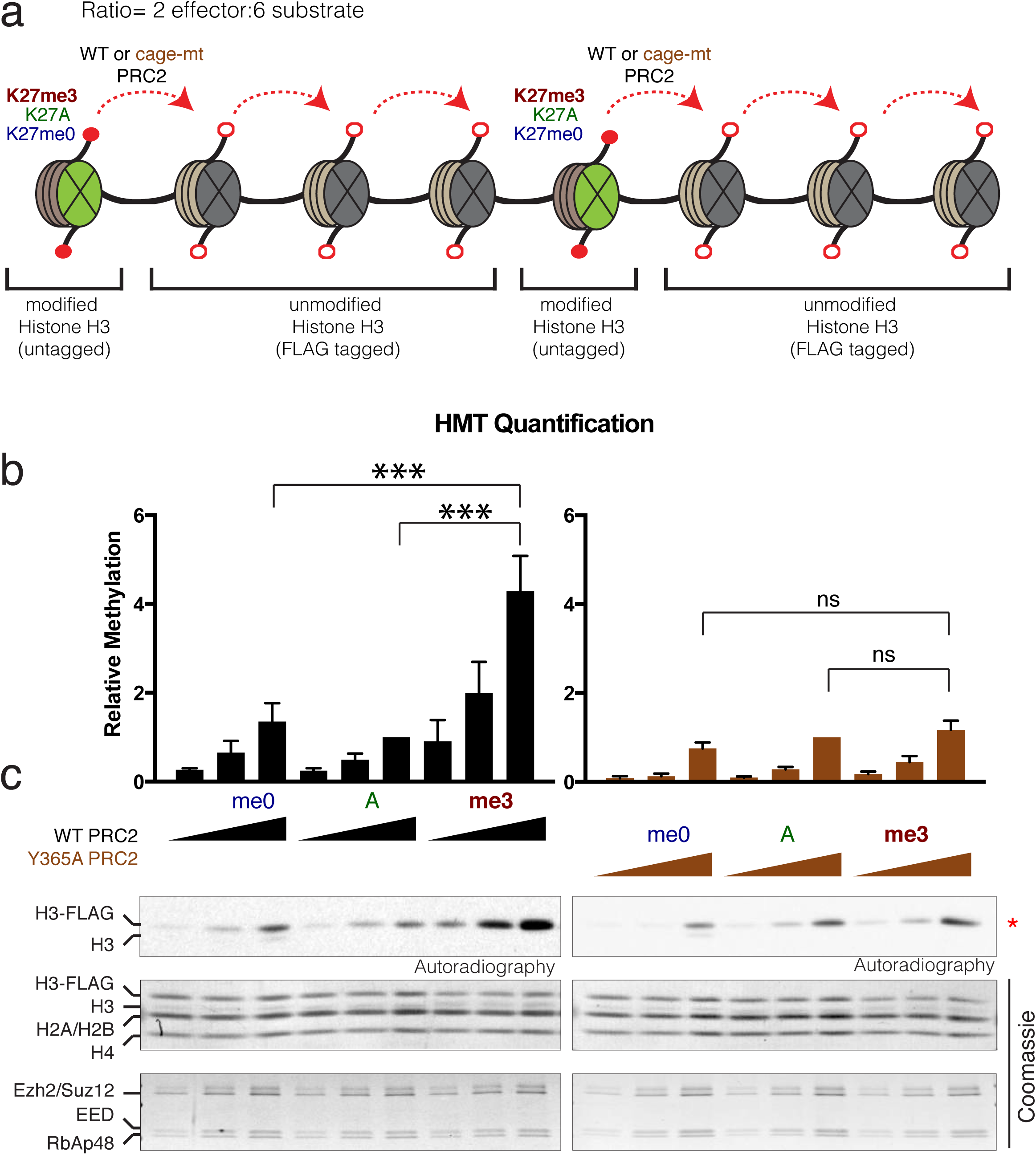
EED-H3K27me3 interaction stimulates PRC2 activity in cis. **a,** Design of the *in vitro* histone methytransferase assay (HMT). The oligonucleosome arrays comprised K27me0 (WT), K27A or K27me3 modifications on both histone H3 within the effector nucleosome (green) and FLAG-tagged unmodified histone H3 within the substrate nucleosome (gray), at a ratio of 2 effector:6 substrate nucleosomes. Recombinant PRC2 reconstituted with either WT or cage-mutant EED (Y365A) was gauged for activity towards substrate nucleosomes. **b**, Quantification of the methylation levels scored by autoradiography (below) on the unmodified FLAG-tagged H3 with indicated concentrations of PRC2 (n=3, one sided paired test: ***p-value<0.005). **c**, Extent of methylation obtained the *in vitro* HMT assay, as scored by autoradiography. Total levels of histones and PRC2 components as shown by Comasssie blue staining.

**Figure S7:**
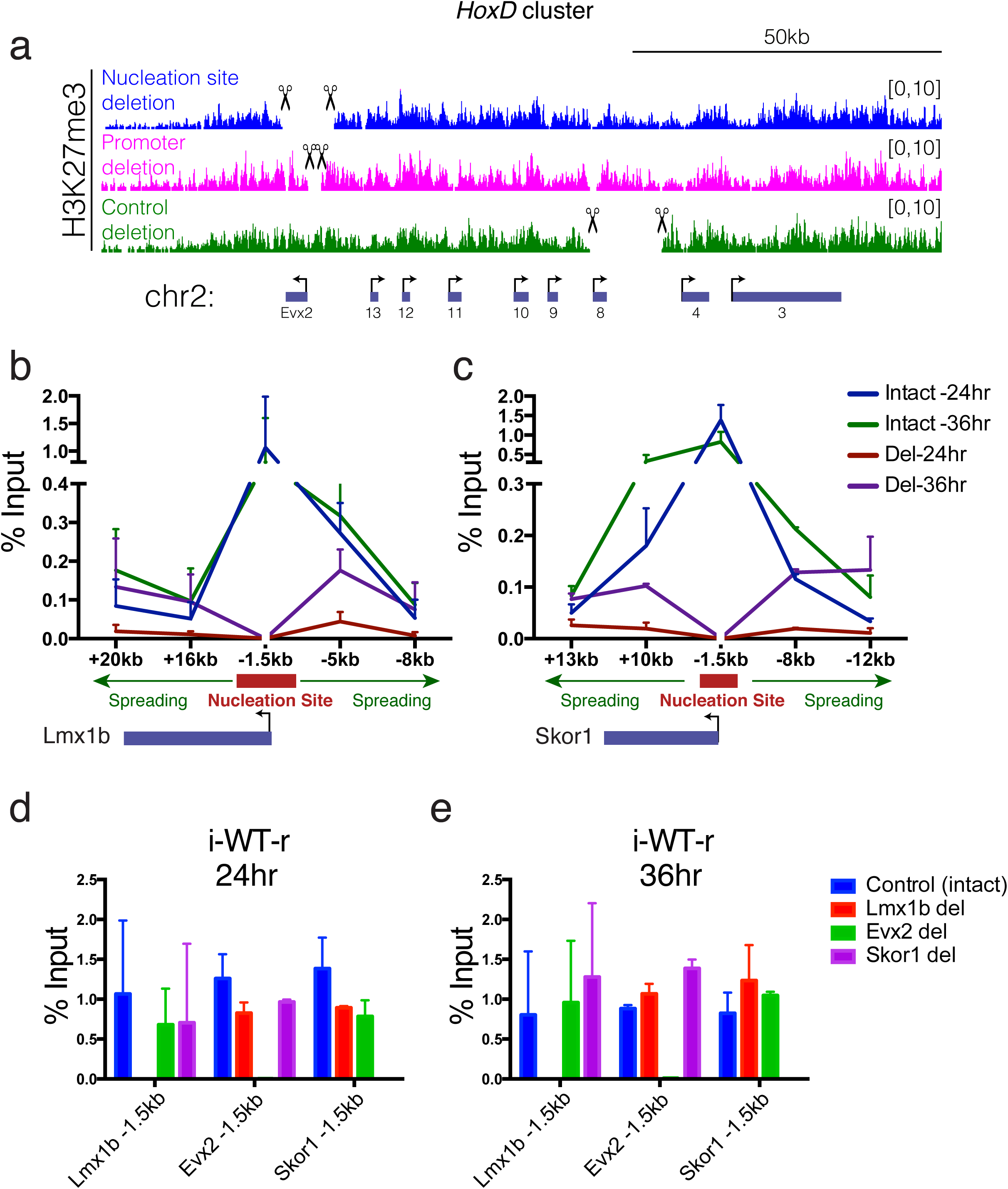
Deletion of a nucleation site delays H3K27me3 deposition to spreading sites. **a**, ChIP-seq tracks for H3K27me3 along the *HoxD* cluster in mESCs harboring the depicted genomic deletions in steady-state mESCs. **b**, **c**, H3K27me3 ChIP-qPCR comparing intact versus deleted nucleation sites at 24 hr and 36 hr after addition of 4-OHT in i-WT-r cells, at the indicated regions relative to the TSS of *Lmx1b* (**b)**, and *Skor1* **(c)**. Enrichments were calculated as % of input. **d**, **e**, H3K27me3 ChIP-qPCR using the indicated primers at 24 hr **(d)** and 36 hr **(e)** of WT rescue in mESCs with the indicated nucleation site deletions. Enrichments were calculated as % of input. All ChIP-qPCR experiments were performed in two biological replicates.

**Figure S8:**
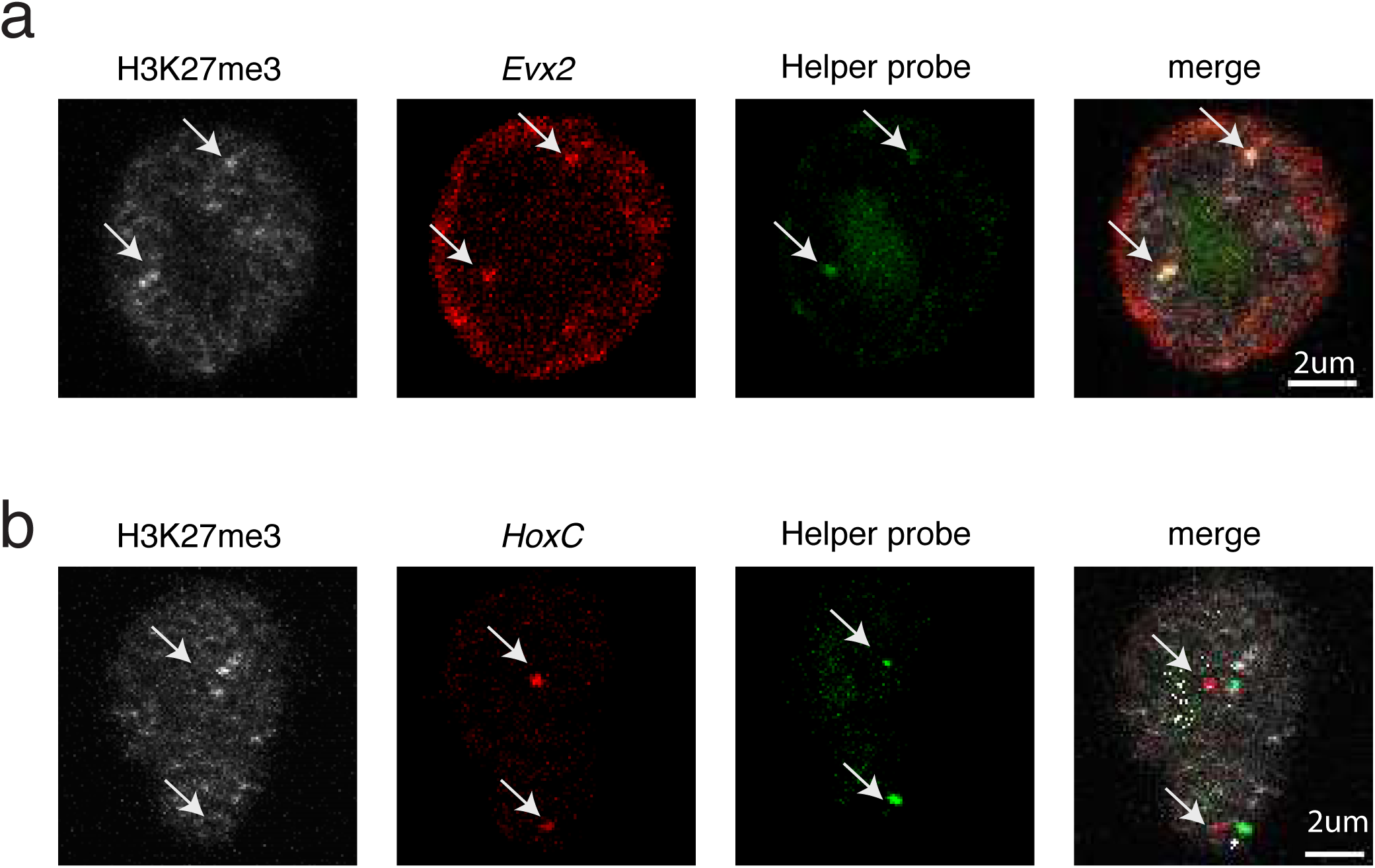
Nucleation site overlaps with H3K27me3 foci. **a, b,** ImmunoFISH using H3K27me3 antibody, a nucleation site probe (*Evx2*) **(a)** or a spreading site probe (*HoxC*) **(b)** at 12 hr after WT EED expression in i-WT-r cells. A helper probe, which is specific to a region in the vicinity of *Evx2* or *HoxC,* was used to determine the real FISH signal. Arrows indicate the localization of the nucleation site and spreading site alleles. Images are examples of 2 biological replicates that are quantified in Figure S5b.

**Figure S9:**
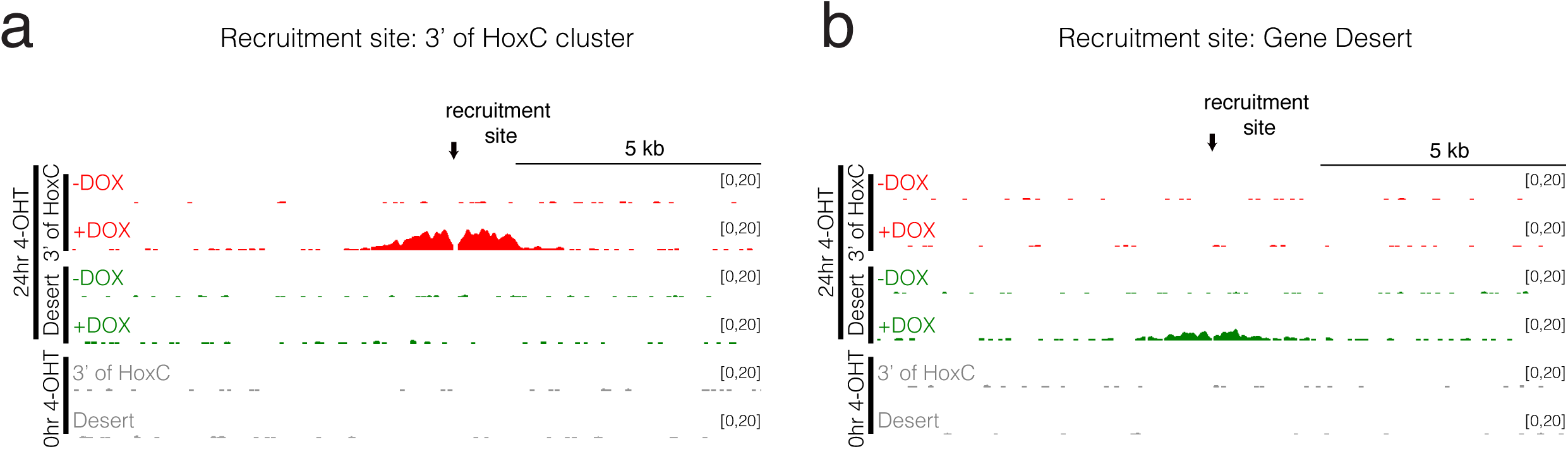
Artificial recruitment of TetR-EZH2 to the 3‘ end of the *HoxC* cluster or a gene desert results in formation of ∼5 kb of H3K27me3 domains. **a**, **b**, ChIP-seq tracks for H3K27me3 showing deposition of H3K27me3 after induction of TetR-EZH2 recruitment to 3‘ of *HoxC* **(a)** and a gene desert **(b)** upon DOX treatment in i-WT-r cells after WT EED expression (24 hr, 4-OHT induction). As controls, H3K27me3 levels in the absence of WT EED expression (0 hr, without 4-OHT) at each recruitment site are indicated.

**Figure S10:**
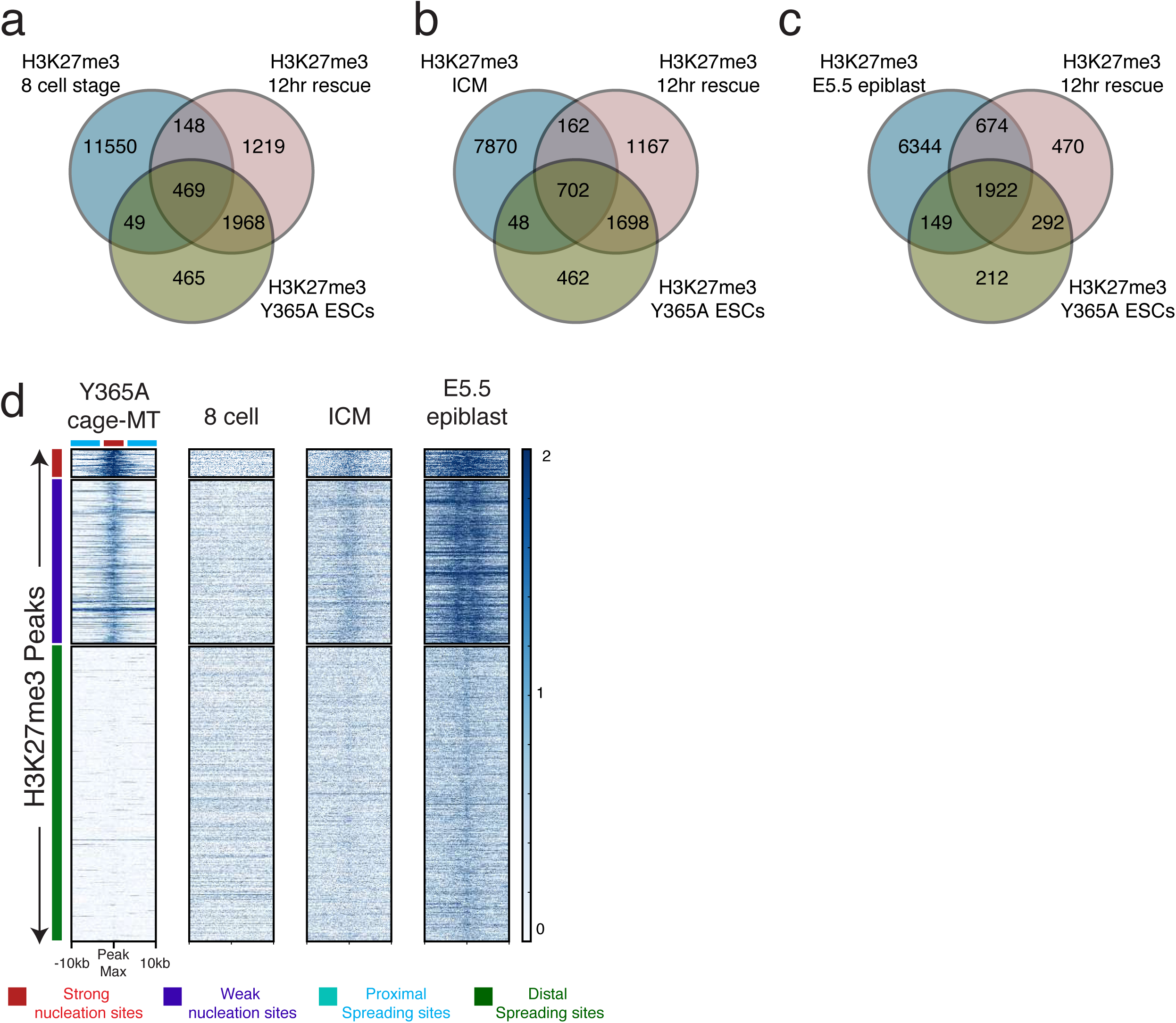
*De novo* deposition of H3K27me3 to nucleation sites occurred as early as the ICM stage during mouse development. **a, b, c,** Venn diagram showing the overlap in H3K27me3 peaks between the 12 hr i-WT-r, Y365A from this study and the 8-cell stage **(a)** ICM **(b)** and E5.5 epiblast **(c)** from Zheng *et al*., 2016. Significance of overlaps were determined by Fisher‘s exact test (two-tail); p-values: 12 hr vs. 8-cell= 5.0942e-24; 12 hr vs ICM= 0; 12 hr vs. E5.5 epiblast= 0; Y365A vs. 8-cell= 3.091e-31; Y365A vs. ICM= 0; Y365A vs. E5.5 epiblast= 0. **d,** Heatmaps of H3K27me3 ChIP-seq in the EED Y365A cage-mutant and the indicated developmental stages within a 20 kb window centered on the maximum value of peak signal in WT mESCs. H3K27me3 peaks were sorted in descending order by signal intensity at the 12 hr time-point. Strong nucleation sites are marked in red (total=237), weak nucleation sites in blue (total=1389), proximal spreading sites in cyan and distal spreading sites in green (total= 36618, representative of randomly selected 2500 peaks are indicated). (Scale: 0-2 for H3K27me3).

## Supplemental Information Table Legends

**1: Global % histone modification levels remained largely constant in EED cage-mutants.** Histone modification levels (highlighted in yellow) within the indicated genotypes of EED were identified by quantitative histone mass-spectrometry. The experiments were performed in 3 replicates of which replicate #1 and #2 are technical and #3 is biological. Only 1 replicate of Y365A was performed.

**2: Identification of proteins enriched at chromatin comprising EED, either cage-mutant (Y365A) or WT.** Peak integration values of all peptides identified by ChIP-MS derived from proteins isolated by FLAG pull-down of FLAG-HA tagged EED. Average peak integration values of 3 technical replicates were calculated and normalized to histones in each sample. The normalized values for individual proteins from the EED cage-mutant chromatin were divided by those from WT chromatin (MT/WT). WT mESCs without FLAG-HA tagged protein were also subjected to FLAG pull-down and used as negative control (Neg).

**3: Nucleation sites are in spatial proximity to each other in WT EED mESCs.** Enrichment analysis of nucleation sites on 4C domains in WT EED. Significant overlaps of nucleation site with 4C domains over a background generated by random shuffling of the 4C domains are determined using ‘fisher’ function in bedtools. Fisher‘s exact test was applied to determine the significant enrichments of nucleation sites within the 4C domains. 4C experiments were performed in 2 biological replicates.

**4: Nucleation site overlaps with H3K27me3 foci.** Quantification of overlap between H3K27me3 foci at 12 hr and a nucleation site *(Evx2)* or a spreading site *(HoxC)* by ImmunoFISH. Colocalization was defined by an overlap of at least two pixels between the signals from the DNA BAC probe and the H3K27me3 antibody. Two independent biological replicates are shown. Fisher‘s exact test was applied to determine the statistical difference between *Evx2* and *HoxC.*

**5:** List of CRISPR constructs

**6:** List of Antibodies

**7:** List of ChIP-seq experiments

**8:** List of ChIP-qPCR primers

**9:** List of RNA-seq experiments

**10:** List of 4C-seq primers

## Methods

### Mouse ESC culture and differentiation

E14Tga2 (ATCC, CRL-1821) and C57BL/6 (gift from Agnel Sfeir) ESCs were grown in standard medium supplemented with LIF, 1 µM MEK1/2 inhibitor (PD0325901, Stemgent) and 3 µM GSK3 inhibitor (CHIR99021, Stemgent). For motor neuron differentiation, the protocol in Narendra et al. was used (Narendra et al., 2015). For neural progenitor differentiation, 300K ESCs were plated in suspension plates with medium containing 50% Advanced DMEM/F12 (Life Technologies: 12634-028), 50% Neurobasal Medium (Life Technologies: 21103-049), 15% Knockout Serum Replacement (Life Technologies: 10828-028), 1% Pen/Strep, 2 mM L-Glutamine, and 0.1 mM 2-mercaptoethanol to generate embryoid bodies (EBs). 4 days later, the EBs were treated with 5 µM all-trans-Retinoic acid (RA, Sigma) for 4 days to generate neuronal progenitor cells. For cardiac differentiation, the protocol in Takahashi et al. was used (Takahashi et al., 2003).

### CRISPR genome editing

gRNAs were designed using CRISPR design tool in https://benchling.com. All gRNAs in Table S5 were cloned in pSpCas9(BB)-2A-GFP (PX458, a gift from Feng Zhang, Addgene plasmid #48138) and transfected into ESCs either with oligo donor (ODN) or donor DNA (Table S5), cloned in PCR Blunt vector (Life Technologies) using Lipofectamine 2000 (Life Technologies). Single clones from GFP positive cells were genotyped and confirmed by sequencing.

### Generation of inducible WT or cage-mutant EED rescue cells (Figure 2a)

First, exon 10 and 11 of the endogenous copy of EED was deleted in mESCs possessing a CreERT2 transgene. Then a cassette within the intron following exon 9 of EED was introduced, which comprised its remaining 3‘ cDNA sequence and a C-terminal Flag-HA tag upstream of T2A-GFP, all in reverse orientation with respect to the endogenous gene sequence. The cassette was flanked by a splice-acceptor and polyadenylation sequence nested between heterologous inverted loxP sites (lox66 and lox71) (Zhang and Lutz, 2002). Upon tamoxifen administration (4-OHT), exon 9 spliced directly into the cassette, producing an epitope-tagged WT or cage-mutant EED protein along with a T2A-GFP marker to identify cells in which recombination was successful. All of these genome editing studies were performed using gRNAs and donor DNAs in Table S5.

### Immunofluorescence

ESCs were fixed with 4% paraformaldehyde in PBS and permeabilized with PBS/0.25% Triton X-100 at RT for 30 min. After blocking with blocking buffer (PBS/5% donkey serum/0.1% Triton X-100) at RT for 30 min, primary antibodies were incubated overnight at 4°C. Several washes were performed with washing buffer (PBS/0.1% Triton X-100), before secondary antibody incubation. After several more washes and DAPI staining, cells were mounted with Aqua mount (Ref 13800) and imaged with Zeiss Airyscan 880 confocal microscopy at 63X magnification. Images are processed and pseudo-colored using a distribution of ImageJ, Fiji (Schindelin et al., 2012).

### Immunoprecipitation

Nuclear extracts were prepared as in Gang Li et al. (Li et al., 2010) Following overnight antibody incubation, agarose beads were added and incubated for 2 hr at 4°C. After washing with BC200 with 0.1% NP40, antibody-antigen interactions were eluted with 2X Lamelli Buffer and subjected to western blot.

### Chromatin immunoprecipitation followed by mass spectrometry (ChIP-MS)

Nuclei were extracted using HMSD buffer (20mM HEPES pH 7.5 at 4°C, 5mM MgCl2, 250mM sucrose, 1mM DTT). Nuclei were resuspended in Mnase buffer (20mM HEPES pH 7.5 at 4°C, 100mM NaCl, 5mM CaCl2). The nuclei were treated with limited amount of Mnase at 37°C for 10 min, adjusted to yield nuclosomal lengths that average from 2-7 nucleosomes. After the reaction was terminated with equal volume of buffer with 10mM EGTA, the nuclei were ruptured by brief sonication to liberate the nucleosomal chains. Supernatant was collected and incubated with Flag beads to pull down Flag-HA tagged cage-mutant (Y365A) or WT EED (Sigma) for 4hr at 4°C. Next, the beads were washed with BC50 (20mM HEPES pH 7.5 at 4°C, 50mM NaCl, 5mM MgCl2) and elution was done with Flag peptide in BC50 (Sigma). All buffers include protease and phosphatase inhibitors as well 5mM sodium butyrate. Eluted samples were trypsinized and subjected to standard data-dependent LC-MS/MS over a two-hour reverse phase gradient. Peptides were identified by Mascot and summed peptide peak integration values for individual proteins were acquired using Skyline Software.

### ChIP-seq

ChIP-seq experiments were performed as described previously (Gao et al., 2012). In brief, cells were fixed with 1% Formaldehyde. Nuclei was isolated using LB1 (50 mM HEPES, pH 7.5 at 4°C, 140 mM NaCl, 1 mM EDTA, 10% Glycerol, 0.5% NP40, 0.25% Triton X; 10 min at 4°C), LB2 (10 mM Tris, pH 8 at 4°C, 200mM NaCl, 1 mM EDTA, 0.5 mM EGTA; 10 min at RT) and LB3 (10 mM Tris, pH 7.5 at 4°C), 1 mM EDTA, 0.5 mM EGTA, 0.5% N-Lauroylsarcosine sodium salt) buffers in order. Chromatin was fragmented to an average size of 250 bp using a Diagenode Bioruptor. Chromatin immunoprecipitation was performed with the antibodies listed in Table S6. Chromatin from *Drosophila* (in a 1:50 ratio to the ESC-derived chromatin) as well as *Drosophila* specific H2Av antibody was used as spike-in control in each sample. For ChIP-seq, libraries were prepared as described in Narendra et al. (Narendra et al., 2015) using 1-30 ng of immunoprecipitated DNA. All ChIP-seq experiments are listed in Table S7. ChIP-qPCRs were performed with 2X SYBR Green Master PCR mix (Roche), and detected by Stratagene Mx3005p instrument. All ChIP-qPCR primers are listed in Table S8.

### RNA-seq

Total RNA from ESCs and EBs (day 2) was isolated with TRIzol (Life Technologies) and reverse transcribed using Superscript III and random hexamers (Life Technologies) to synthesize the 1st strand. Second strand was synthesized with dUTP to generate strand asymmetry using DNA Pol I (NEB, M0209L) and the E. coli ligase (Enzymatics, L6090L). RNA-seq libraries were constructed using the protocol described in Narendra et al. (Narendra et al., 2015). All RNA-seq experiments are listed in Table S9.

### 4C-seq

The protocol from van de Werken et al. (van de Werken et al., 2012) was used to prepare 4C-seq libraries. HindIII-DpnII inverse primers for the bait regions were chosen from https://compgenomics.weizmann.ac.il/tanay/?page_id=367. Illumina compatible 5′ adapter overhangs from van de Werken et al. (van de Werken et al., 2012) was included in the primers for high-throughput sequencing. All 4C-seq primers are listed in Table S10.

### Data analysis

#### a) ChIP-seq

Sequence reads for ChIP-seq were mapped with Bowtie 2 using default parameters (Langmead et al., 2009). After normalization with the spike-in *Drosophila* read counts, ChIP-seq densities were visualized on the USCS genome browser (https://genome.ucsc.edu/). Heatmaps for the ChIP-seq experiments were generated using deepTools (Ramirez et al., 2014). Average ChIP-seq read density profiles were plotted using R programming. Venn diagrams to visualize the extend of the overlap among ChIP-seq samples were drawn using ‘ChIPpeakAnno’ package from Bioconductor (Zhu et al., 2010).

#### b) RNA-seq

RNA-seq experiments were analyzed as previously reported (Narendra et al., 2015). Briefly, sequence reads were mapped with Bowtie (Langmead et al., 2009) and normalized differential gene expression was calculated with DEseq (R package) (Anders and Huber, 2010).

#### c) 4C-Seq

Sequence reads from 4C-Seq were aligned and processed as described before using 4C-ker (Raviram et al., 2016). The cisAnalysis function was used to call interacting 4C domains for each viewpoint. For the HoxB, Specc1 and Pou5f1 baits a k of 5 was used and for the rest k of 10 was used. 4C-tracks were visualized by IGV (Robinson et al., 2011). To compare the number of overlapping nucleation sites with 4C interacting domains, a two tailed fishers exact test in ‘fisher’ function of bedtools was used (Quinlan and Hall, 2010).

### Liquid Chromatography and Quantitative Histone Mass Spectrometry

Total nuclear histones were isolated by standard acid extraction (Lin and Garcia, 2012). The histones were next derivatized with propionic anhydride, which adds propionyl groups to unmodified and monomethylated lysines prior to trypsinization. Post-trypsinization, the peptides were derivatized with propionic anhydride which adds a propionyl group to the newly formed n-termini. Analytical columns were produced in-house by pulling fused silica microcapillary tubing (75um i.d.) with a flame to generate a tip. The columns were packed with ReproSil-Pur C18-AQ resin (3um, Dr. Maisch GmbH, Germany) using a pressurized cylinder. A Thermo Easy NanoLC 1000 HPLC (Thermo Scientific, Odense, Denmark) was used to load sample (1.5ug) onto the column. Samples were resuspended in 0.1% formic acid prior to loading. Peptides were then separated using reverse-phase chromatography composed of two buffers: buffer A: 0.1% formic acid in water, buffer B: 0.1% formic acid in acetonitrile. A 60-minute gradient was applied to the column at 300nL/min flow rate: 0-28%B in 45 minutes, 28-80%B in 5 minutes, 80%B for 10 minutes. Samples were eluted into a hybrid linear ion trap-Orbitrap (Orbitrap Elite, Thermo Scientific, Bremen, Germany). Full MS scans (300-1100 m/z) were collected in the Orbitrap at a resolution of 120,000 and an AGC target of 5 × 105. MS/MS scans were obtained using collision induced dissociation (CID) with a normalized collision energy of 35 in the ion trap with an AGC target of 3 × 104. The method first collected a full MS spectra followed by 8 MS/MS spectra covering 300 – 700 m/z in 50 m/z windows (i.e. 300-350 m/z, 350-400 m/z, etc.). A second full MS was obtained followed by 8 more MS/MS spectra covering 700-1100 m/z in 50 m/z windows. Mass spectrometry data were analyzed using EpiProfile software (Yuan et al., 2015) and the quants for the H3 (27-40aa) peptides were further verified manually.

### Protein expression and purification

6xHIS-tagged EZH2, FLAG-tagged EED, Suz12, and RbAp48 were subcloned in pFASTBac1 baculovirus expression plasmid. Recombinant PRC2 core complex was produced in SF9 cells grown in SF-900 III SFM (Invitrogen). After 60 h of infection, SF9 cells were harvested and resuspended in BC150 (25 mM Tris-HCl, pH 7.9, 0.2 mM EDTA, 150 mM KCl, 10% glycerol) with 0.1% NP40 and protease inhibitors (1 mM phenylmethlysulfonyl fluoride (PMSF), 0.1 mM benzamidine, 1.25 mg/ml leupeptin and 0.625 mg/ml pepstatin A). Cells were then sonicated and PRC2 was purified through Ni-NTA agarose bead (Qiagen), FLAG-M2 agarose beads (Sigma), and Q Sepharose column (GE healthcare).

### Hetero-oligonucleosome preparation

Methylated histones were generated by using the Methyl Lysine Analog (MLA) strategy as described previously (Simon et al., 2007; Simon and Shokat, 2012). Octamers were prepared as described previously (Lusser and Kadonaga, 2004; Yun et al., 2012). To generate hetero-oligonucleosomes, indicated ratio of effector and substrate octamers were mixed with 8x 601 sequence containing linearized DNA templates and subjected to salt dialysis (Lusser and Kadonaga, 2004; Yun et al., 2012). The concentrations of hetero-oligonucleosomes were determined by quantifying the amount of nucleosomal DNA on 0.8 % agarose gel visualized by ethidium bromide staining.

### Histone methyltransferase (HMT) assay

Standard HMT assays were performed as described previously (Margueron et al., 2009; Son et al., 2013). Briefly, the reaction was performed in a total volume of 15 ml of HMT buffer (50 mM Tris-HCl, pH 8.5, 5 mM MgCl_2_, 4 mM DTT) with ^3^H-labeled SAM, substrate (recombinant nucleosomes), and the indicated concentration of PRC2, either WT or comprising EED Y365A. The reaction mixture was incubated for 1 h at 30°C and separated by SDS-PAGE gel electrophoresis. The gel was transferred to a PVDF membrane. The membrane was exposed to X-ray film (Kodak XAR) to detect the level of methylation as analyzed by autoradiography exposure.

### DNA FISH with immunofluorescence

Bacterial artificial chromosomes (BACs) containing *Exv2* (RP23-331E7, RP23-364J22), *Hoxc13* (RP23-430C12), and a nearby probe to help locate *Hoxc13* (RP24-119A11) were chosen using USCS browser and were ordered from BACPAC (http://bacpac.chori.org/). These BACs were labeled by nick translation with 5-(3-aminoallyl)-dUTP (Life Technologies) then converted into the required fluorophore with Alexa Fluor 647 and Alexa Fluor 555 Reactive Dye (Life Technologies). For each 22×22 mm coverslip, 1ug of labeled BAC DNA, 1µg of sheared salmon sperm DNA (Ambion) and 1µg of Cot-1 (Life Technologies), 1ug Mouse Hybloc DNA (Applied Genetics Lab) were precipitated and resuspended in 20µl of hybridization buffer (50% formamide/20% dextran sulfate/5X denhardt‘s solution). The fluorescence *in situ* hybridization (FISH) protocol was adapted from Chaumeil et al., 2008(Chaumeil et al., 2008). Briefly, 1×106 cells were adhered to poly-L-lysine coated coverslips, fixed for 10min with 2% (wt/vol) paraformaldehyde/PBS on ice, washed three times in cold PBS, and permeabilized for 10min on ice with 0.5% (vol/vol) Triton/PBS. Samples were then washed three times in cold PBS, then incubated for 1hr at 22°C in blocking solution (2.5% (wt/vol) BSA, 10% (vol/vol) normal goat serum and 0.1% (vol/vol) Tween-20 in PBS). Samples were incubated for 1hr at 22°C with a rabbit antibody raised against H3K27me3 (Cell Signalling technology) diluted 1:200 in blocking solution. Cells were washed three times with 0.2% (wt/vol) BSA / 0.1% (vol/vol) Tween-20/PBS then incubated for 1hr at 22°C with donkey anti-rabbit IgG Alexa Fluor 488 (Thermofisher) diluted 1:500 in blocking solution. Cells were washed three times with 0.1% (vol/vol) Tween-20 in PBS. Cells were post-fixed for 10min at 22°C in 2% (wt/vol) paraformaldehyde, briefly rinsed in 2X saline sodium citrate (SSC), then incubated with RNaseA (0.1 mg/ml in 2XSSC) for 1hr at 37°C. The cells were then re-permeabilized for 10min on ice in 0.7% (vol/vol) Triton-X-100 / 0.1 M HCl. Probes and cells were then denatured together on glass slides for 2min at 75°C. Slides were then incubated overnight at 37°C. Coverslips were then removed from the slides, and washed three times with 50% (vol/vol) formamide, 2XSSC at 42°C, followed by three washes in 2XSSC at 42°C, and three washes in 2XSSC at 22°C. Coverslips were then mounted in Prolong Gold (Life Technologies) containing 1.5µg/ml 4,6-diamidino-2-phenylindole (DAPI, Sigma).

### Confocal microscopy and analysis

DNA FISH with immunofluorescence was imaged by confocal microscopy on a Leica SP5 AOBS system (Acousto-Optical Beam Splitter). Optical sections separated by 0.3 µm were collected and only cells with H3K27me3 signal were analyzed using ImageJ software (NIH). Signal for probes that were scored (RP23-331E7 for *Exv2* and RP23-430C12 for *Hoxc13*) were confirmed using localization of “helper” probes (RP23-364J22 for *Evx2* and RP24-119A11 for *Hoxc13*), which appear in a limited nuclear volume near the scored probe. Probes were defined as co-localized with H3K27me3 if the DNA probe signals and immunofluorescence foci directly overlapped (at least two pixels of colocalization). Statistical significances were calculated by a two-sided Fisher‘s exact test P-values ≤ 5.00e-2 were taken to be significant. As detailed in each case in the figure legends, P-values displayed in the main figures were applied to combined data from repeated experiments. Error bars represent the standard deviation between experiments.

